# Structural basis for regulation of Frizzled-4 signaling by the co-receptor Tetraspanin-12

**DOI:** 10.1101/2025.09.25.678640

**Authors:** Alyana J. Granados, Payal P. Pratap, Kevin Alexander Estrada Alamo, Katherine J. Susa

## Abstract

Norrin is an atypical ligand that regulates retinal angiogenesis through the Wnt/β-catenin pathway. Norrin triggers heterodimerization of Frizzled-4 (FZD4) with Low-density lipoprotein receptor-related protein 5 or 6 (LRP5/6), leading to downstream β-catenin stabilization. Unlike Wnt ligands, Norrin requires the tetraspanin Tspan12 for signaling amplification, but it is not understood why. Here, we report a 3.4-angstrom structure of Tspan12 in complex with FZD4 determined by cryo-electron microscopy. The structure reveals that FZD4 and Tspan12 form a direct complex in the absence of Norrin. The transmembrane (TM) domain of Tspan12 is oriented in a tightly packed four-helix bundle and interacts with TM2 of FZD4 to promote trafficking of Tspan12 to the cell surface. The C-D helices of Tspan12, which mediate binding to Norrin, remain exposed while Tspan12 is in complex with FZD4, facilitating higher-affinity Norrin binding. Cell-based assays reveal that Tspan12 and FZD4 remain associated after Norrin recognition, suggesting that Tspan12 is a core component of the FZD4-Norrin-LRP5/6 signaling complex. Together, these studies reveal the molecular mechanism underlying the assembly of the Tspan12-FZD4 complex and the enhancement of Norrin signaling, suggesting new ways to target the FZD4-Tspan12 complex for ocular diseases characterized by hypo- or hyper-vascularization.

**SIGNIFICANCE STATEMENT:** Blood vessel formation in the retina is essential for vision, and its disruption leads to ocular diseases. Norrin regulates retinal angiogenesis by triggering the heterodimerization of Frizzled-4 with co-receptor LRP5/6, mimicking the function of Wnt ligands and leading to downstream β-catenin signaling. Unlike Wnts, Norrin requires a tetraspanin family member, Tspan12, to amplify signaling. We report the structure of the FZD4-Tspan12 co-receptor complex and identify key interacting regions that enable Tspan12 trafficking to the cell surface and higher affinity capture of Norrin. Cell-based assays indicate Tspan12 remains a component of the active Norrin-FZD4-LRP5/6 signaling complex. Our results highlight an underappreciated role of tetraspanins in direct regulation of receptor signaling and suggest new therapeutic avenues for targeting abnormalities in vascular growth.

## INTRODUCTION

Wnt/β-catenin signaling is a highly conserved cellular communication pathway that is one of the main regulators of development, cell fate determination, tissue homeostasis, and tissue regeneration in metazoans, (1). Wnts, a family of secreted, lipid-modified growth factors, initiate signaling by binding to G-protein coupled receptors (GPCRs) of the Frizzled family, which contain an extracellular N-terminal cysteine-rich domain (CRD) (2, 3). Wnt binding within the CRD (4, 5) promotes heterodimer formation with the co-receptor Low-density lipoprotein receptor-related protein 5 or 6 (LRP5/6)(6). FZD-LRP5/6 heterodimers recruit the intracellular signaling effectors Dishevelled (Dvl) (7) and Axin (8), ultimately leading to stabilization of the transcriptional co-activator β-catenin and induction of Wnt target genes to drive cell division, migration, and differentiation (9). In line with this, dysregulated signaling can lead to developmental defects, cancer, and degenerative disorders (1, 10).

In the early 1990s, an atypical ligand that activates β-catenin signaling, Norrin (Norrie Disease Protein, NDP), was discovered through experiments aimed at identifying genes associated with Norrie disease, a neurodevelopmental disorder characterized by severe vision loss or blindness, intellectual disability, and hearing loss (11, 12). Norrin has a transforming growth factor-β-like cysteine knot fold and is homodimeric, consisting of two disulfide-linked protomers (13). Norrin has no structural homology to Wnts yet mimics their function. Each Norrin protomer contains non-overlapping binding sites for the CRD of FZD4 and for LRP5/6, which triggers heterodimerization of FZD with LRP5/6 to initiate β-catenin signaling (14). Unlike Wnt ligands, which bind promiscuously to multiple FZD family members (15), Norrin specifically binds to FZD4 out of the ten human Frizzled receptors (16). Norrin/FZD4 signaling is essential for retinal blood vessel formation and blood-retina barrier development (17). Consistent with this critical role, mutations in genes encoding components of this pathway—including Norrin, FZD4, and LRP5—cause familial exudative vitreoretinopathy (FEVR), a hereditary ocular disorder related to Norrie disease characterized by retinal hypo-vascularization, leading to retinal detachment, vision loss, or blindness (18, 19). Additionally, Norrin plays important roles in angiogenesis not only for eye development but also in the ear, brain, and female reproductive system (13).

Compared to Wnt ligands, Norrin has the unique feature of utilizing a member of the tetraspanin family as a co-receptor to amplify β-catenin signaling. Nearly two decades after the identification of Norrin, a large-scale reverse genetic screen revealed that mutations in the gene encoding for Tetraspanin-12 (Tspan12) mimic the retinal vascular defects of NDP, FZD4, and LRP5 mutant mice (20).Tspan12 is a member of the ancient, highly conserved tetraspanin family of four-pass transmembrane proteins, which have been termed “molecular facilitators” (21). Tetraspanins bind transmembrane partner proteins to regulate a range of fundamental signal transduction pathways (22, 23), including B cell (24, 25), Notch (26), and integrin (27, 28) signaling through regulation of partner protein trafficking and membrane microdomain localization. Cell-based assays demonstrated that Tspan12 functions as a co-receptor to significantly increase Norrin/β-catenin signaling by about 4-fold, with no effect on Wnt/β-catenin signaling (20). In line with Tspan12’s important role in amplifying Norrin signaling, mutations in Tspan12 cause FEVR and account for approximately 5-10% of FEVR cases in humans (18, 19, 29–33).

The mechanism underlying Tspan12’s enhancement of FZD4 signaling has remained unclear, with proposed models including the allosteric modulation of FZD4 to amplify signaling (34), the promotion of FZD4 clustering (35), or the enhancement of FZD4/LRP5 heterodimerization (36). Tspan12 can rescue β-catenin signaling disrupted by point mutations that impair the Norrin-FZD4 interaction, indicating that Tspan12 may enhance the binding of Norrin to FZD4 (34). However, it is unclear whether Tspan12 achieves this function through a direct interaction with FZD4. Recent biophysical experiments showed that Tspan12 directly binds Norrin with high affinity (10 nM), and can bind to the same protomer of Norrin as the CRD of FZD4 (37). However, Tspan12 is competitive with LRP6 for binding to the same site on Norrin (37).

These data support a ligand hand-off mechanism, in which Tspan12 initially promotes the capture of Norrin and then passes it off to FZD4 and LRP5/6 (37). An alternative mechanism is Tspan12 remaining incorporated in the Norrin-FZD4-LRP5/6 complex, with Tspan12 and LRP5/6 each binding to separate protomers of Norrin. However, the structural basis for Tspan12 engaging a single protomer remains unclear.

To understand how Tspan12 mediates enhancement of Norrin signaling, we determined the structure of Tspan12 bound to FZD4 using single-particle cryo-electron microscopy (cryo-EM). We then used cell-based signaling assays to characterize how Norrin binds to the Tspan12-FZD4 complex and monitor the dynamics of Tspan12-FZD4 upon Norrin recognition. Our structural analysis and cell-based assays define the molecular basis for Tspan12 enhancement of Norrin signaling and suggest a model in which Tspan12 does not “hand off” Norrin to FZD4 and LRP5 but instead remains associated as a constitutive component of the FZD4-LRP5/6 signaling complex.

## RESULTS

### Engineering and structure determination of the Tspan12-FZD4 complex

Tspan12 requires interaction with FZD4 for trafficking to the cell surface (34) and co-localizes with FZD4 on the cell surface (13, 34, 35), suggesting a direct interaction that initiates in the endoplasmic reticulum (34). However, our initial attempts to purify a stable Tspan12-FZD4 complex through co-expression and dual-affinity tag purification were unsuccessful, likely due to weak affinity between Tspan12-FZD4, which is consistent with prior reports for both Tspan12-FZD4 and other tetraspanin-partner complexes (37, 38). To overcome this, we engineered a construct with Tspan12 tethered to FZD4 to increase their local concentration and apparent affinity. This construct contains the full-length FZD4 sequence connected to full-length Tspan12 with a flexible glycine-glycine-serine (GGS) linker **(Supplementary Figure 1A)**.

To validate that this tethered construct maintains the natural interaction that facilitates FZD4-mediated trafficking of Tspan12 to the cell surface, we used a flow cytometry-based trafficking assay to monitor Tspan12 surface expression with an anti-Tspan12 antibody **(Figure 1A)**. While Tspan12 expressed alone showed low levels on the cell surface, co-expression with FZD4 resulted in a 4-fold increase in Tspan12 trafficking to the cell surface **(Figure 1A)**. The tethered Tspan12-FZD4 construct displayed a similar magnitude of increase in Tspan12 surface trafficking as Tspan12 co-transfected with FZD4, indicating that our engineered construct retains the interaction that mediates Tspan12 trafficking and is properly folded **(Figure 1A)**.

**Figure 1:**
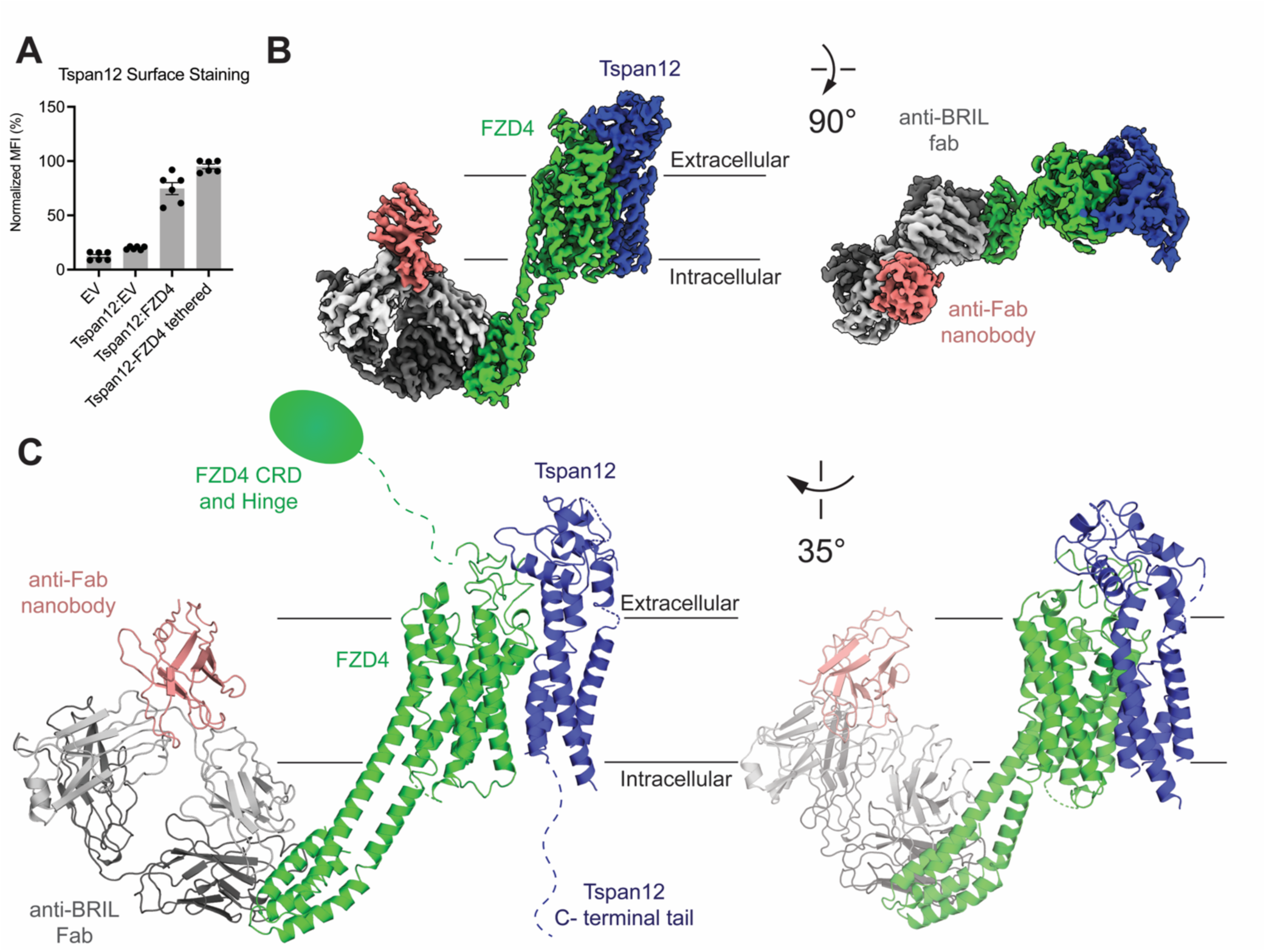
**3D reconstruction and atomic model of the human FZD4-Tspan12 complex bound to an anti-BRIL F_ab_ and anti-F_ab_ nanobody (PDB: 9YEV).** (**A**) Tspan12 trafficking assay with Tspan12 either transfected with empty vector (EV), FZD4, or as a tethered complex with FZD4. Surface Tspan12 was detected by flow cytometry using an Alexa 647-coupled anti-Tspan12 antibody (Biotechne FAB8910R-025). (**B**) Different views of the cryo-EM map of the FZD4-Tspan12-anti-BRIL F_ab_-anti-F_ab_ nanobody complex, colored by subunit. (**C**) Ribbon representation of the structure of the complex. FZD4, Tspan12, the anti-F_ab_ nanobody, and the anti-BRIL F_ab_ are colored as in B. In Panel A, error bars represent mean ± SEM of two independent experiments, with three technical replicates each.

Initial structure determination efforts for the Tspan12-FZD4 complex by single particle cryo-EM were unsuccessful due to the relatively small size of the complex (93 kDa) and limited rigid extracellular features outside of the detergent micelle. To overcome this, we introduced the thermostabilized BRIL domain (cytochrome b652 RIL) into intracellular loop 3 (ICL3) of FZD4, following previously reported successful structure determination of inactive state GPCRs that used an anti-BRIL Fab (BAG2) as a rigid fiducial marker to improve particle alignments (39, 40). To optimize a rigid connection between FZD4 and BRIL, we used AlphaFold3 to guide linker selection, choosing a linker that was predicted to form a continuous rigid helix with high confidence **(Supplementary Figure 1B-D)** (41). We confirmed that the tethered FZD4-BRIL-Tspan12 construct maintained the ability to traffic Tspan12 to the cell surface. Our BRIL-engineered construct resulted in enhanced levels of Tspan12 and FZD4 on the cell surface **(Supplementary Figure 1E, F)**, indicating that the BRIL domain likely stabilizes FZD4, leading to higher expression levels, which is consistent with prior reports (42).

For structure determination, we incubated purified FZD4-BRIL-Tspan12 with anti-BRIL Fab (43) and an anti-Fab nanobody (44) **(Supplementary Figure 1G, H)**. Cryo-EM imaging of this complex revealed well-dispersed particles, and two-dimensional class averages showed clear secondary structures in both the Fab region and transmembrane region **(Supplementary Figure 2)**. Subsequent 3D refinement resulted in a structure of FZD4-BRIL-Tspan12 bound to the anti-BRIL F_ab_ and anti-F_ab_ nanobody at an overall nominal resolution of 3.4 Å, with density well defined for all features of the complex, except the N-terminal CRD-Hinge region of FZD4 and the C-terminal tail of Tspan12, both of which are highly flexible **(Figure 1B, Supplementary Table 1)**.

### Tspan12 does not alter FZD4 conformation

To build the initial model, AlphaFold3 (45) models of FZD4-BRIL and Tspan12, along with the structures of the anti-BRIL-F_ab_ and anti-F_ab_ nanobodies (39), were docked into the cryo-EM density map and manually rebuilt. While the FZD4 model fit readily into the density map, the transmembrane helices of Tspan12 had to be separately docked due to differences in their orientation compared to the AlphaFold3 model **(Supplementary Figure 3)**. In the complex, Tspan12 packs against the second and third transmembrane domains (TM2 and TM3) of FZD4 via hydrophobic interactions and also makes contacts with extracellular loop 1 (ECL1) and the lower region of the hinge domain (also termed “linker”) of FZD4, which is a flexible region connecting the CRD to the 7-transmembrane domain (40, 46) **(Figure 1C, Supplementary Figure 3B-D)**. The FZD4-Tspan12 complex buries a total of approximately 1338 Å^2^ at the interface between the ectodomains and transmembrane domains.

One model of Tspan12 enhancement of signaling is that Tspan12 allosterically activates FZD4 to enhance binding of the intracellular signaling effector Dvl (34). However, recent experiments have shown that the affinity of the Dvl2 DEP domain for the Tspan12-FZD4 complex is not significantly different than its affinity for Fzd4 alone (37). In line with this, the FZD4-Tspan12 structure shows no change in FZD4 conformation compared to the inactive state crystal structure of FZD4 (PDB: 6BD4), with a root mean square deviation (RMSD) of 0.68 Å **(Supplementary Figure 4A)**, indicating that Tspan12 is not functioning to allosterically activate FZD4 (47). A structure of FZD4 bound to the DEP domain of Dvl2 was also determined by cryo-EM (PDB: 8WMA), which revealed that DEP induces a subtle outward movement of TM6 (4.0 Å) and a slight inward movement of TM5 of FZD4 (48). Tspan12 binds to TM2 and TM3 of FZD4 and makes no contacts with FZD4 that would interfere with DEP engagement, indicating that Tspan12 can remain bound to FZD4 after Dvl engagement **(Supplementary Figure 4B)**. While it is possible that the BRIL fusion may influence these structural comparisons, our results indicate that it is unlikely that Tspan12 plays a significant allosteric role.

### The C-D helices of Tspan12 are exposed within the FZD4 complex to facilitate higher affinity Norrin binding

Tspan12 is the only known tetraspanin identified to date with an obvious receptor function for an endogenous secreted ligand, binding to Norrin within its ectodomain with an affinity of approximately 10 nM (34, 37, 49). In line with this, Tspan12 has a particularly unique mode of engagement with FZD4 compared to other tetraspanin-partner structures solved to date **(Figure 2A and 2B)**. In addition to their four-pass transmembrane domain, tetraspanins have two extracellular domains, a small extracellular loop (SEL) and a large extracellular loop (LEL) (23). Within the LEL, tetraspanins have two hypervariable helices, the “C” and “D” helices. Among different members of the tetraspanin family, these helices are highly variable in sequence and structure, representing an important site where structural variability in tetraspanins encodes specificity for their interactions with partners (23). Two other structures of tetraspanins with non-tetraspanin partners have been reported to date at a resolution sufficient to build an atomic model—the tetraspanin CD81 with the B cell co-receptor CD19, and Tspan15 with the metalloprotease ADAM10 (38, 50). In both of these structures, the C and D helices form the primary interaction interface with the partner protein. However, in the structure of Tspan12 bound to FZD4, the C and D helices are completely exposed and highly dynamic, making up the area of poorest local resolution in the cryo-EM map **(Supplementary Figure 5).** The C and D helices were recently shown to be the site of Norrin binding to Tspan12 using a high-confidence AlphaFold-Multimer prediction and model-guided point mutagenesis (37). We used a flow-cytometry-based binding assay to measure the EC_50_ of Norrin binding to the FZD4-Tspan12 tethered construct with an E170K mutation in Helix D of Tspan12, which is predicted by the Norrin-Tspan12 AlphaFold model to form a hydrogen bond with Serine 83 of Norrin (37). This construct bound Norrin with a 2-fold decrease in apparent affinity, confirming the C and D helices of Tspan12 mediate binding to Norrin within the context of the FZD4-Tspan12 complex **(Figure 2C)**.

**Figure 2:**
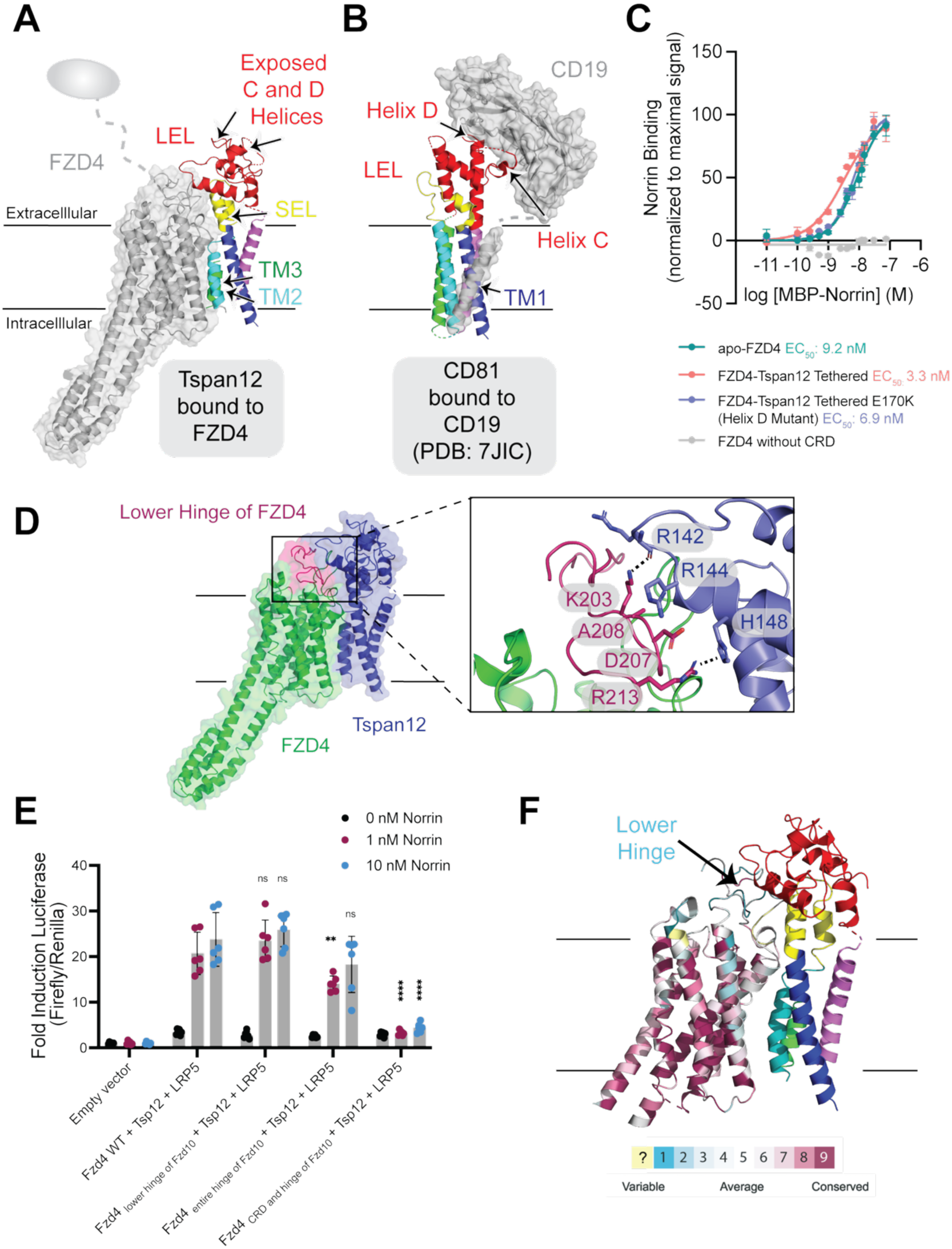
Characterization of Norrin binding to the FZD4-Tspan12 complex. (A and. **B)** Comparison of the FZD4-Tspan12 complex to the CD19-CD81 (Tspan28) complex (PDB: 7JIC). The partner protein is colored in grey, and the tetraspanin is colored by domain. The tetraspanin CD81 utilizes the C-D helices within the LEL as the primary engagement site with its partner, CD19. In contrast, the structure of FZD4-Tspan12 shows that these two helices do not make contact with the partner. **(C)** Binding of Norrin to Expi293 cells expressing either apo-FZD4, tethered FZD4-Tspan12, tethered FZD4-Tspan12 with an E170K point mutation in the D helix, or FZD4 without the cysteine-rich domain (CRD) (negative control). Apo-FZD4 EC_50_: 9.2 nM (95% CI: 7.3 nM to 12.2 nM), Hill slope: 1.0 (95% CI: 0.9 to 1.2). Tethered FZD4-Tspan12 EC_50_: 3.3 nM (95% CI: 2.0 nM to 6.5 nM), Hill slope: 0.8 (95% CI: 0.6 to 1.1). Tethered FZD4-Tspan12 E170K EC_50_: 6.9 nM (95% CI: 6.0 nM to 8.4 nM), Hill slope: 1.1 (95% CI: 1.0 to 1.3). Experiments were performed in triplicate with error bars representing mean ± SEM. In all binding experiments, FZD4 without the CRD was used as a control for background binding. **(D)** Interaction between the lower hinge of FZD4 and Tspan12, showing a zoomed-in interface. **(E)** TOPFlash signaling assay with FZD4/FZD10 chimeras with swaps in the lower hinge or entire hinge domain, or with the entire CRD and hinge (negative control). Expression of chimeric FZD constructs is shown in Supplementary Figure 6. Error bars represent mean ± SEM of two independent experiments, with three technical replicates each. Statistical analysis was performed in GraphPad Prism using an unpaired t-test, comparing wild-type FZD4:Tspan12:LRP5 to each corresponding Norrin concentration of the chimeric FZD4:Tspan12:LRP5 conditions. *p ≤ 0.05, **p ≤0.01, ***p ≤0.001, ****p ≤0.0001, ns: not significant. **(F)** Structure of FZD4-Tspan12, with FZD4 colored by residue conservation among the 9 human Frizzled paralogs. Residues are colored on a sliding scale from magenta (highly conserved) to blue (poorly conserved). Residues with insufficient information for analysis are colored yellow. Conservation was determined using the Consurf server (51).

Tspan12 and FZD4 bind to non-overlapping regions on the same protomer of Norrin (37). Because the C and D helices of Tspan12 remain accessible for Norrin capture when Tspan12 is bound to FZD4, we hypothesized that this may provide avidity to promote increased Norrin capture by positioning the CRD (the Norrin binding domain of FZD4) and the C-D helices of Tspan12 in close proximity. To test this, we measured the EC_50_ of Norrin binding to the tethered FZD4-Tspan12 construct compared to FZD4 alone using a flow cytometry-based binding assay. The tethered construct bound Norrin with approximately 3-fold higher apparent affinity, with an EC_50_ of 3.3 nM (95% confidence interval (CI): 2.0 nM to 6.5 nM) for the FZD4-Tspan12 complex and 9.1 nM (95% CI: 7.3 nM to 12.2 nM) for apo-FZD4 **(Figure 2C)**. These data indicate that one function of Tspan12 is to enable response to low Norrin concentrations.

### Tspan12 interacts with the lower hinge of FZD4

The Tspan12-FZD4 structure revealed that the LEL of Tspan12 interacts with the lower region of the hinge domain of FZD4 **(Figure 2D, Supplementary Figure 3B-C**). The hinge domain connects the CRD to the 7-TM domain of FZD receptors. This domain is highly variable in length and sequence among FZD family members and can be subdivided into a highly flexible segment from the CRD to the first conserved cysteine residue, followed by a shorter, more ordered segment from the first conserved cysteine to TM1 (40). While the CRD and upper hinge segment are not observed in our structure due to inherent flexibility, a portion of the lower, ordered hinge segment makes several contacts with the LEL of Tspan12. Within the hinge, Lysine 203 of FZD4 hydrogen bonds with the carbonyl of Arginine 142 of Tspan12, and Arginine 213 of FZD4 interacts with Histidine 148 of Tspan12 **(Figure 2D, Supplementary Figure 3B-3C)**.

**Figure 3:**
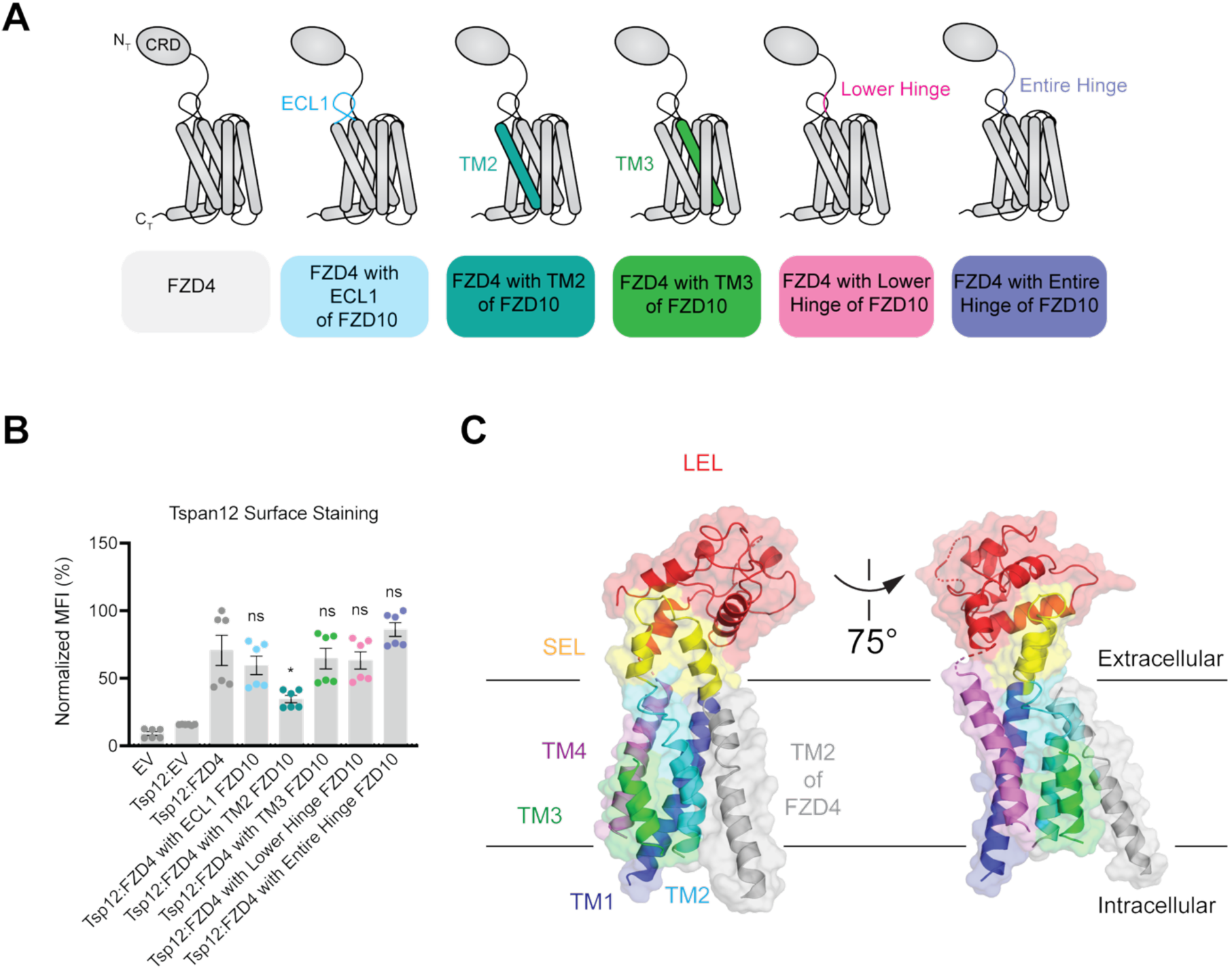
**The second transmembrane domain of FZD4 is essential for Tspan12 trafficking to the cell surface**. **(A)** Schematic of FZD4-FZD10 chimeras. ECL1, TM2, TM3, or the hinge of FZD4 was replaced with that of FZD10. **(B)** Tspan12 trafficking assay with Tspan12 co-transfected with empty vector (EV), FZD4, or a FZD4-FZD10 chimera. Surface Tspan12 was detected by flow cytometry using an Alexa 647-coupled anti-Tspan12 antibody (Biotechne FAB8910R-025). Error bars represent mean ± SEM of two independent experiments, with three technical replicates each. Statistical analysis was performed in GraphPad Prism using an unpaired t-test, comparing Tspan12:FZD4 to each Tspan12: chimeric FZD4 condition. *p ≤ 0.05, **p ≤0.01, ***p ≤0.001, **** p≤0.0001. Expression of chimeric FZD constructs is shown in Supplementary Figure 7. **(C)** Atomic representation of Tspan12 bound to FZD4, colored by domain. TM2 of FZD4 (grey) packs against TM2, TM3, and the SEL of Tspan12.

The hinge domain of FZD4 has been demonstrated to play a role in signaling. Swapping the FZD4 hinge with the FZD5 hinge results in a loss of Norrin signaling and a decrease in affinity for Norrin (52). We next asked if Tspan12’s interaction with the lower hinge region of FZD4 contributes to Norrin signaling. To test this, we created FZD4 chimeras in which either the entire hinge domain or just the lower hinge of FZD4 is replaced with that of FZD10, and a negative control chimera in which the entire CRD and hinge were replaced with that of FZD10, and assessed their ability to signal in the TOPFlash β-catenin reporter assay **(Supplementary Table 2, Supplementary Figure 6A)**. As expected, the FZD4 chimera with the CRD and hinge swap does not signal, and the FZD4 chimera with the entire hinge region swapped with that of FZD10 showed decreased signaling at low Norrin concentrations **(Figure 2E)**. However, the FZD4 chimera with the lower hinge swapped for that of FZD10 signaled comparably to wild-type FZD4 at all Norrin concentrations tested **(Figure 2D)**. The lower hinge domain is highly sequence divergent among the ten human Frizzled receptors, suggesting that, though this region does not have an apparent role in signaling, it may help drive the specificity of the interaction of Tspan12 with only FZD4 out of the ten receptors **(Figure 2F)**.

### Interactions within the transmembrane domain mediate Tspan12 trafficking to the cell surface

Tspan12 and FZD4 initiate their partnership in the secretory pathway, and Tspan12 is dependent on FZD4 for proper trafficking to the cell surface (34). This is highly unusual for a member of the tetraspanin family—in other tetraspanin-partner trafficking relationships studied to date, the tetraspanin is responsible for promoting the trafficking of the partner to the cell surface (53). For example, the tetraspanin CD81 promotes the trafficking of its partner, CD19, to the cell surface by covering an exposed, highly hydrophobic patch within the CD19 ectodomain, thereby enhancing the surface expression of properly folded CD19, with CD19 having no reciprocal effect on CD81 expression (38, 54).

Next, we asked how FZD4 promotes trafficking of Tspan12 to the cell surface. We created a panel of FZD4 chimeras with domain swaps in domains that interact with Tspan12 **(Figure 3A, Supplementary Table 2, Supplementary Figure 7)**. We replaced either ECL1, TM2, TM3, the lower hinge, or the entire hinge of FZD4 with that of FZD10 and used a flow cytometry-based trafficking assay to monitor the ability of these constructs to support Tspan12 surface expression. While Tspan12 expressed without FZD4 showed low levels on the cell surface, co-expression of Tspan12 with FZD4 resulted in a 4-fold enhancement of Tspan12 expression on the cell surface **(Figure 3B)**. While the ECL1, TM3, and hinge chimeras had no effect on Tspan12 trafficking, replacement of TM2 with that of FZD10 significantly reduced the amount of Tspan12 on the cell surface, indicating that the interaction of Tspan12 with TM2 of FZD4 is a determinant of trafficking **(Figure 3B).** In addition to Tspan12 abundance on the plasma membrane, blotting whole cell lysate of cells co-transfected with Tspan12 and either wild type FZD4 or chimeric constructs confirmed that in the presence of the TM2 chimera, significantly less Tspan12 is present in the cell, indicating that Tspan12 is either being made less efficiently or is rapidly degraded in the presence of the TM2 swap chimera **(Supplementary Figure 8).** TM2 of FZD4 packs against the SEL and TM2 and TM3 of Tspan12, stabilizing a highly bent region between TM2 and the SEL of Tspan12 (**Figure 3C, Supplementary Figure 3C**). Tspan12 can bind to Norrin with high affinity in the absence of FZD4 (37), but it cannot signal unless it is complexed with FZD4 (35). This unusual trafficking relationship for a tetraspanin may have evolved to prevent apo-Tspan12 from being expressed in high amounts on the cell surface. Apo-Tspan12 would be able to bind to Norrin, but when not complexed to FZD4, it could act as a “sink” for Norrin by binding ligand but not being in close enough proximity to FZD4 for efficient signaling.

### Dynamics of the FZD4-Tspan12 interaction

A “hand-off” mechanism for Tspan12 function was proposed in which Tspan12 initially promotes the capture of Norrin and then passes off Norrin to FZD4 and LRP5/6, leading to downstream signaling (37). This model is based on evidence that Tspan12 and LRP6 compete for the same binding site on Norrin (37), suggesting that Tspan12 may dissociate from FZD4 after Norrin recognition to facilitate LRP5/6 heterodimerization. We and others have previously shown that tetraspanins are conformationally dynamic, switching between an “open” state (38) and a “closed” state (55). The “open” conformation of the tetraspanin CD81 has a higher affinity for its partner CD19 (38, 55, 56), and we hypothesized that conformational changes in Tspan12 between an open and closed state may facilitate dissociation from FZD4. While the “closed” conformation has a cone-shaped orientation of the transmembrane helices and a large hydrophobic binding pocket capable of binding lipids (55, 57), the “open” conformation has a tightly packed four-helix bundle orientation of the TMs, which occludes the lipid binding pocket, and is characterized by a hinge opening of the ectodomain away from the membrane (38, 58). When bound to FZD4, Tspan12 is in the “open” conformation, with a tightly packed orientation of the TMs, similar to the “open” conformation of CD81. **(Figure 4A, 4B)**.

**Figure 4:**
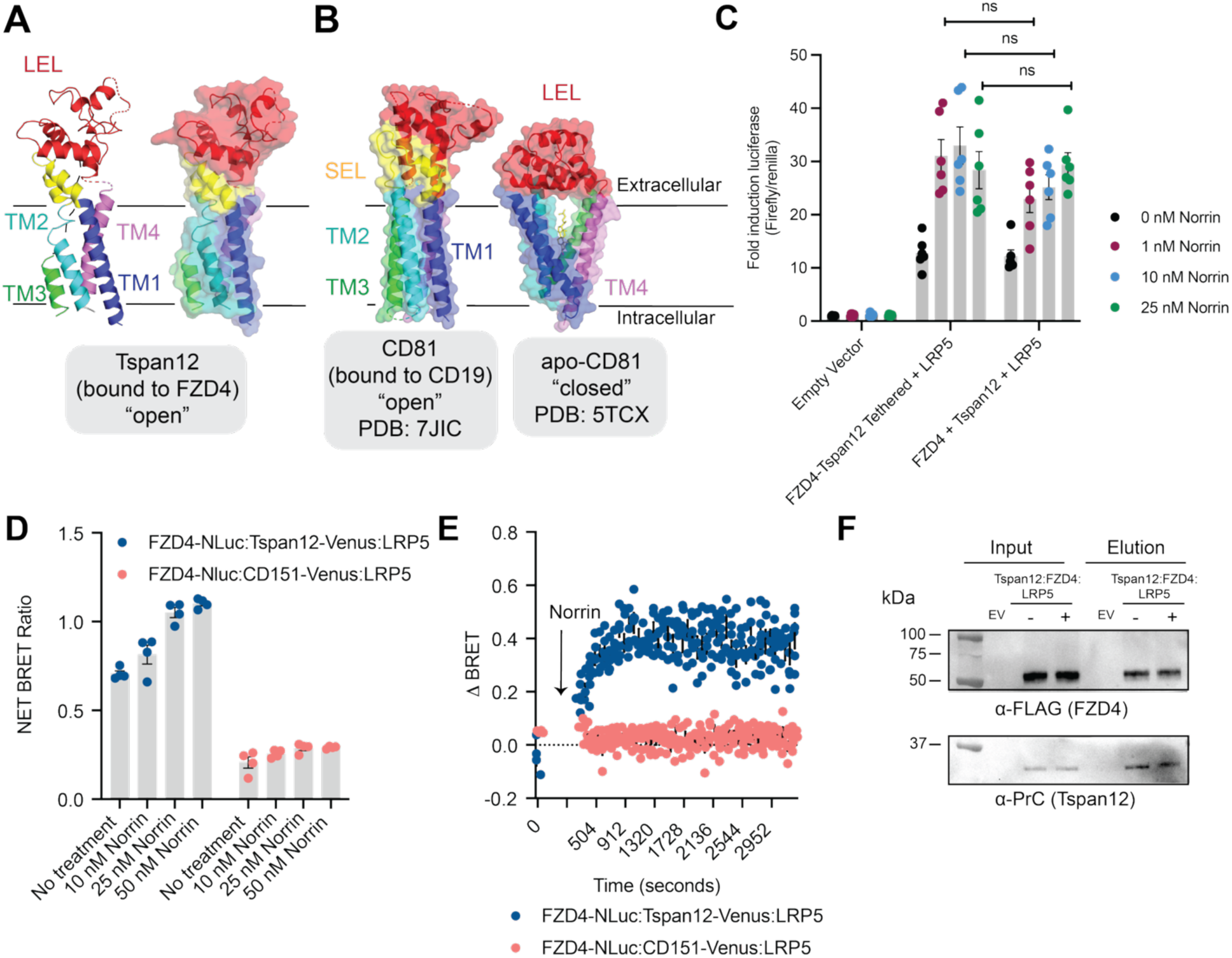
Tspan12 remains as a core component of the FZD4 signaling complex after Norrin recognition. **(A)** Structure of Tspan12 (bound to FZD4), highlighting an “open” state of Tspan12, with a closely packed four helix bundle orientation of the transmembrane helices. Tspan12 is colored by domain. **(B)** Comparison of “closed” apo-CD81 (right, PDB: 5TCX) with “open” CD19-complexed CD81 (left, PDB: 7JIC). The structures are colored by domain. The “open” conformation refers to an opening of the LEL away from the membrane. **(C)** TOPFlash signaling assay with tethered FZD4-Tspan12 or co-transfected FZD4 and Tspan12. Error bars represent mean ± SEM of two independent experiments, with three technical replicates each. Statistical analysis was performed in GraphPad Prism using an unpaired t-test, comparing tethered and co-transfected FZD4 and Tspan12 at each corresponding Norrin concentration. *p ≤ 0.05, **p ≤0.01, ***p ≤0.001, ns: not significant. Expression levels of FZD4 and Tspan12 constructs and co-transfection signaling controls are shown in Supplementary Figure 6. **(D)** BRET assay tracking Tspan12-FZD4 proximity. Fixed amounts of LRP5 and FZD4-NLuc were transfected into ΔFZD_1–10_ knockout cells with either CD151-Venus or Tspan12-Venus to assess the change in Tspan12-FZD4 proximity after 5 minutes of treatment with either 10 nM, 25 nM, or 50 nM Norrin. **(E)** Kinetic monitoring of Tspan12-FZD4 proximity upon 10 nM Norrin treatment in ΔFZD_1–10_ knockout cells expressing LRP5 and FZD4-NLuc with either Tspan12-Venus or the negative control, CD151-Venus. For BRET experiments in Panels D and E, data are presented as mean ± SEM of five technical replicates **(F)** Pulldown of PrC tagged Tspan12 associated with FLAG tagged FZD4 in untreated cells (-) or after 10 nM Norrin treatment (+) for 15 minutes in ΔFZD_1–10_ knockout cells co-transfected with PrC-Tspan12, FLAG-FZD4, and untagged LRP5. EV: empty vector transfected cells. The blot in panel F represents one biological replicate that is representative of two biological replicates.

To investigate whether Tspan12 dissociation from FZD4 is required for signaling, we utilized our tethered FZD4-Tspan12 construct to determine if forcing the association of Tspan12 with FZD4 via covalent tethering inhibits signaling. If Tspan12 dissociation from FZD4, allowing for LRP5/6 engagement, is a key step in the signaling mechanism, we would expect that the forced recruitment of Tspan12 to FZD4 via tethering would result in decreased signaling, thereby preventing LRP5/6 engagement and downstream signaling. Using the TOPFlash β-catenin reporter assay in ΔFZD_1–10_ cells, we compared the ability of the tethered construct or co-transfected FZD4 and Tspan12 to induce signaling under conditions with matched receptor levels (**Figure 4C, Supplementary Figure 6B**). Tethered Tspan12-FZD4 signaled at levels matching those of co-transfected FZD4 and Tspan12, indicating that dissociation of Tspan12 from FZD4 is not strictly required for signaling.

In a series of orthogonal assays, we used a bioluminescence resonance energy transfer (BRET) assay to monitor the association of Tspan12 with FZD4 in cells. We fused the luciferase energy donor NanoLuc (NLuc) to FZD4 and energy acceptor Venus to Tspan12, or a negative control, non-interacting tetraspanin, CD151, using previously reported linkers and insertion sites **(Supplementary Table 2)** (59, 60). When in close proximity (within 10 nm (61)), NLuc and Venus undergo BRET, generating a fluorescent signal. We used this system to determine if the amount of Tspan12 in proximity to FZD4 changes after Norrin recognition. In cells co-transfected with LRP5 and the BRET constructs, we either left the cells untreated or treated them with 10, 25, or 50 nM Norrin, and then measured the BRET ratio 5 minutes after Norrin addition **(Figure 4D)**. Importantly, Tspan12-Venus and FZD4-NLuc showed a 4- to 5-fold increase in the BRET ratio compared to our negative control, CD151-Venus, highlighting the assay’s ability to detect the true interaction between FZD4 and Tspan12 **(Figure 4D)**. If Tspan12 is dissociating from FZD4 after Norrin treatment, we would expect to see a decrease in the BRET ratio, indicating decreased proximity. However, the BRET ratio was slightly increased following Norrin treatment, again suggesting that Tspan12 is not dissociating from FZD4 **(Figure 4D)**. To assess if Tspan12-FZD4 proximity changes over the time course of signaling, we transfected ΔFZD_1–10_ cells with our BRET constructs and LRP5, collected a baseline reading, left the cells untreated or added 10 nM Norrin, and then collected a time course of the BRET signal over a period of approximately 55 minutes **(Figure 4E)**. Again, we observed that the BRET ratio was slightly increased and remained constant throughout our entire data collection period, further suggesting that Tspan12 remains a core component of the FZD4 complex after ligand recognition. Moreover, co-immunoprecipitation of Tspan12 and FZD4 in cells with or without Norrin treatment showed similar amounts of Tspan12 pulled down in both conditions, providing additional evidence that Tspan12 remains associated with FZD4 **(Figure 4F)**. Flow cytometry confirmed that most of the transfected receptor was found on the cell surface rather than in the endoplasmic reticulum, confirming that our pulldown result wasn’t confounded by a large pool of intracellular receptor that is insensitive to Norrin binding **(Supplementary Figure 9).** We saw no evidence for Tspan12 dissociation in three orthogonal signaling assays. Indeed, the affinity of Norrin for Tspan12-FZD4 (low nM) is much higher than the measured affinities of Norrin for LRP6 (∼0.4-2.8 µM) (13, 62), suggesting that the efficiency of this proposed competitive hand-off would be very low in cells. Instead, our data indicate that Tspan12 remains a component of the Norrin signaling complex, which is also consistent with a prior report that Tspan12 co-localizes with the FZD4-LRP5 signaling complex after Norrin binding at the cell surface and in endosomes (63).

## DISCUSSION

Here, we report the structure of the co-receptor Tspan12 bound to FZD4, showing that Tspan12 and FZD4 form a direct complex in the absence of Norrin. Tspan12 engagement of FZD4 enables higher affinity binding to Norrin, and our cell-based assays indicate that Tspan12 remains a core component of the FZD4-LRP5/6 signaling complex after Norrin recognition. This raises a fundamental question: if Tspan12 is not dissociating from FZD4, how is LRP5/6 able to be recruited to the complex? One potential mechanism is that Tspan12 remains incorporated in the Norrin-FZD4-LRP5/6 complex, with Tspan12 and LRP5/6 each binding to opposite protomers of Norrin. Although Norrin is a disulfide-linked homodimer that is overall two-fold symmetric, the cysteine knot fold of Norrin results in distinct spatial orientations of each protomer, breaking perfect symmetry **(Figure 5A).** In the second Norrin protomer, the Tspan12 binding site is rotated approximately 90° **(Figure 5B).** The AlphaFold-Multimer model of Norrin bound to two Tspan12 molecules shows that this orientation results in the second Tspan12 molecule being rotated 90° perpendicular to the membrane plane, which would be physiologically incompatible with binding (**Supplementary Figure 10)**. This suggests a possible model in which Tspan12 binds to one protomer of Norrin and LRP5/6 binds to the opposite protomer of Norrin, leading to the formation of a stable Tspan12-FZD4-LRP5/6 signaling complex **(Figure 5C)**. Alternatively, it is also possible that the binding sites of Tspan12 and LRP5/6 are only partially overlapping, allowing LRP5/6 to engage both protomers of Norrin. The mechanism by which LRP5/6 integrates into the complex and the overall stoichiometry of the active signaling complex remain important open questions.

**Figure 5:**
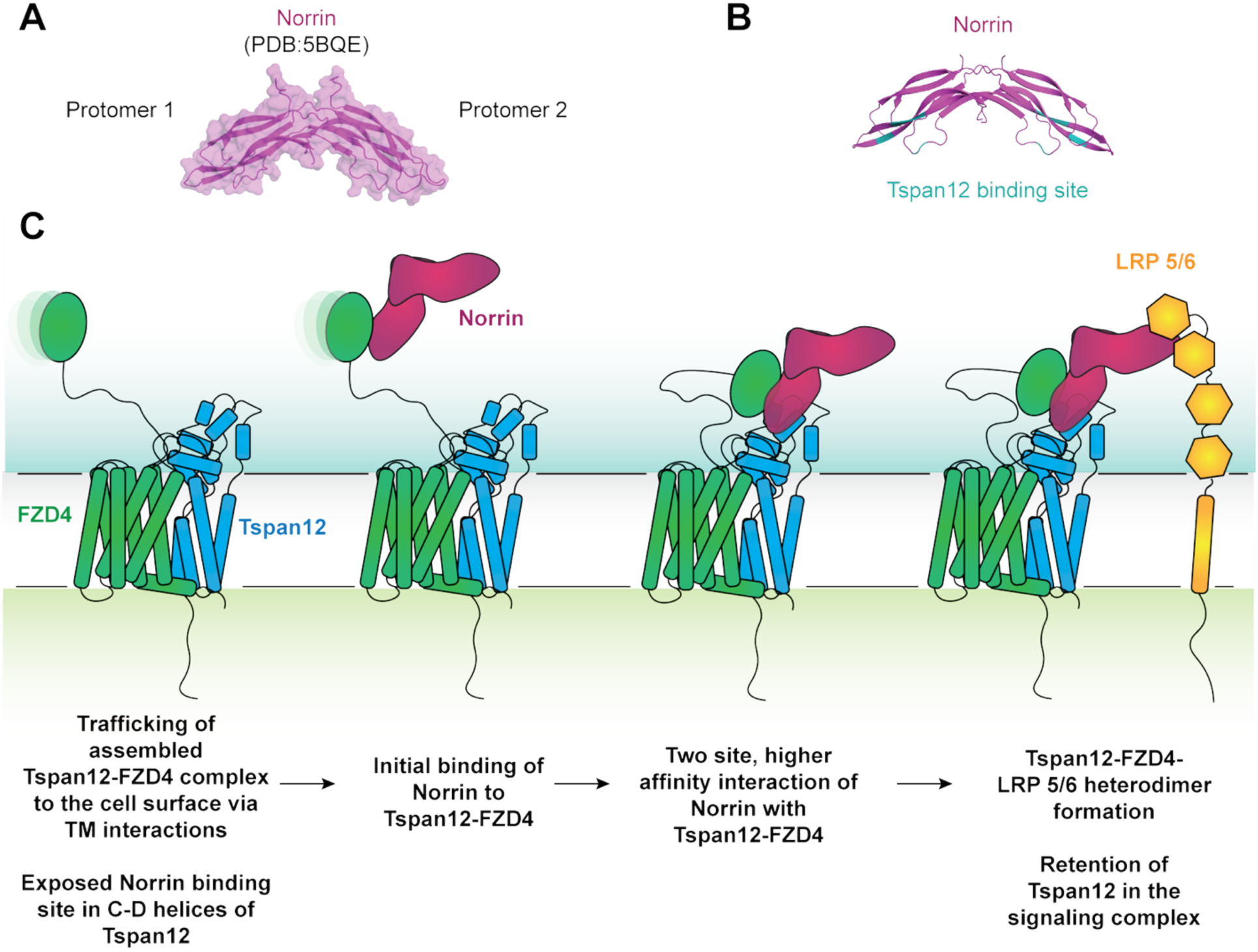
Proposed model for Tspan12 assembly into the Norrin-FZD4-LRP5/6 signaling complex. **(A)** Crystal structure of Norrin (PDB:5BQE). **(B)** Tspan12 binding site(37) mapped onto the surface of Norrin, highlighting distinct spatial orientations of the Tspan12 binding site on each protomer. Residues within 5 Å of Tspan12 in the Tspan12-Norrin AlphaFold3 model are colored in teal. **(C)** Proposed model for incorporation of Tspan12 into the Norrin-FZD4-LRP5/6 signaling complex. Tspan12 is trafficked to the cell surface in conjunction with FZD4 and exists in a complex with FZD4 prior to Norrin recognition. Norrin engages in a two-site interaction with Tspan12-FZD4 to recruit LRP5/6 into a Tspan12-FZD4-LRP5/6-Norrin signaling complex. Although this model depicts LRP5/6 interacting with only one Norrin promoter for simplicity, the mechanism by which LRP5/6 engages Norrin and incorporates into the complex remains an open question.

Wnts contact FZDs through a highly conserved two-site mechanism, positioning “thumb” and “finger” domains on opposite sides of the CRD (5). Why Wnts need a two-site mode of engagement is debated, but one model is that it provides specificity, avidity, and optimally orients the Wnt ligand for LRP5/6 engagement (5, 64). Importantly, Norrin contacts the CRD of FZD4 at a single site (65, 66), suggesting that a function of Tspan12 may provide to additional binding site to facilitate LRP5/6 recruitment and downstream signaling. The C-D helices of Tspan12 are anchored close to the membrane in our structure. Our results suggest that one role of Tspan12 may be to prevent Norrin from diffusing away by providing avidity through its additional binding site and to restrain the motion of Norrin bound to the highly dynamic CRD-Hinge region of FZD4, facilitating proper orientation of Norrin to increase the likelihood of LRP5/6 recruitment. Structural characterization of the native FZD4-Tspan12-LRP5/6-Norrin complex and Wnts in complex with full-length FZD receptors and LRP5/6 will be necessary to answer this question and will remain an important area of future investigation.

The partnership between FZD4 and Tspan12 represents an important, heretofore unappreciated function of tetraspanins in the direct regulation of receptor signaling. Tetraspanins are recognized as “molecular facilitators” (21), acting as chaperones to traffic partner proteins to the cell surface and regulate their plasma membrane localization (22, 53). However, they are not appreciated as a family of membrane proteins that directly signal or bind to endogenous ligands. The finding that Tspan12 directly binds to Norrin with high affinity (37) and our findings that Tspan12 increases the affinity of Norrin binding, while remaining a component of the active Norrin-FZD4-LRP5/6 signaling complex, highlight an important role for tetraspanins in the direct regulation of signaling, rather than simply functioning as molecular chaperones in the membrane. 10 out of 33 human tetraspanins have an intracellular tail, indicating that more tetraspanins than appreciated may play a direct role in signal transduction. The 59 amino acid C-terminal cytoplasmic tail of Tspan12 contains several putative phosphorylation sites, suggesting it may play a role in signaling by scaffolding the recruitment of intracellular signaling molecules. Early work in the tetraspanin field showed that several tetraspanins could co-immunoprecipitate phosphoinositide 4-kinase (67) and protein kinase C (PKC) (68), suggesting that tetraspanins may recruit these kinases to specific membrane locations to influence signaling. More recent work has elegantly shown that in B cells, the N-terminal intracellular domain of tetraspanin CD53 promotes the recruitment of protein kinase C family member PKCβ to CD53-containing microdomains, thereby facilitating PKC-mediated phosphorylation of substrates (69). An important area of future studies will be to characterize whether signaling molecules bind to the intracellular tail of Tspan12 and play a role in amplifying β-catenin signaling, as well as to determine whether any other tetraspanins directly bind an endogenous ligand within their ectodomain.

Modulation of β-catenin signaling is an area of active therapeutic investigation for cancer, degenerative diseases, and diseases related to blood vessel angiogenesis, like FEVR (1, 70). Mutations in Tspan12 are found in approximately 5-10% of FEVR patients (18, 19, 29–33). Mapping known missense mutations found in FEVR patients onto the structure of FZD4-Tspan12 revealed that these mutations do not map to the Tspan12-FZD4 binding interface **(Supplementary Figure 11)**, suggesting instead that these mutations likely decrease Norrin binding or affect protein quality or folding. Targeting the β-catenin pathway remains challenging due to the toxicity associated with broad inhibition of β-catenin signaling. While FZD4 and LRP5/6 are widely expressed throughout the body, Tspan12 has more limited expression (35), offering a potentially powerful way to specifically modulate Norrin signaling for the treatment of diseases of retinal hypo-vascularization, such as FEVR, or for ocular diseases associated with retinal hyper-vascularization, such as diabetic retinopathy. Tspan12 antibodies that inhibit Norrin signaling and decrease angiogenesis have been reported (71). Our results suggest that because Tspan12 forms a direct complex with FZD4, modulates how FZD4 binds to Norrin, and stays engaged with the LRP5/6 signaling complex, antibodies that either block the Norrin binding site on Tspan12 or stabilize the interaction of Norrin with Tspan12 may offer new ways to treat diseases characterized by retinal hyper-vascularization and hypo-vascularization, respectively.

## ACKNOWLEDGEMENTS

This work was funded by an NIH grant DP5OD036136 and a UCSF Sandler Fellowship to K.J.S, and an NIH IMSD training grant 5R25GM056847-25 and Achievement Rewards for College Scientists (ARCS) scholarship to K.A.E.A. We thank members of the Susa and Manglik labs, Dr. Elise Bruguera for helpful discussions, the Manglik lab for the use of equipment, and Dr. Meredith Anne Skiba for the anti-F_ab_ nanobody. We thank Dr. Julia Maria Rogers, Dr. Edward Harvey, and Dr. Aashish Manglik for feedback on the manuscript. Cryo-EM data were collected at the Cryo-EM facility at the University of California, San Francisco. We thank Dr. Glenn Gilbert for helpful comments and assistance during grid screening and data collection. We thank Benoit Vanhollebeke for generating and generously sharing ΔFZD_1–10_ HEK293T cells (72).

## AUTHOR CONTRIBUTIONS

K.J.S. designed and conceived of the project. A.J.G., P.P.P., K.A.E.A, and K.J.S. designed experiments. A.J.G. and K.J.S. prepared the sample for cryo-EM, and A.J.G., K.J.S., and P.P.P. collected and processed cryo-EM data. P.P.P. and K.J.S. performed model building and refinement. A.J.G., P.P.P., K.A.E.A., and K.J.S. performed cell-based assays. K.J.S. wrote the manuscript with input from all authors. K.J.S. supervised the work and provided financial funding.

## DATA AND MATERIALS AVAILABILITY

Materials will be made available upon request. Protein Data Bank (PDB) and Electron Microscopy Data Bank (EMDB) IDs of the FZD4-Tspan12-BRIL F_ab_ model and map are 9YEV and EMD-72864, respectively.

## METHODS

### Design of FZD4-Tspan12 fusion protein

The FZD4-Tspan12 fusion was cloned into a doxycycline-inducible pcDNA3.1 (+) tet/zeo vector with an N-terminal haemagglutinin signal sequence followed by a FLAG epitope tag. Residues 37-537 of human FZD4 (full length, excluding the endogenous signal sequence) were connected to full-length human Tspan-12 (residues 2-305) using a GGGSx4 linker. The linker length was engineered to be long enough in length to ensure tethering of Tspan12 would not enforce Tspan12 engagement on one side of FZD4. For the BRIL FZD4-Tspan12 construct, BRIL replaced ICL3 (residues 416-427) of FZD4 using linkers derived from the A_2A_ adenosine receptor (ARRQL between residue 415 and the N-terminus of BRIL, and ARSTL between the C-terminus of BRIL and residue 428). Tspan12 was linked to FZD4-BRIL with a GGGSx4GGSx3 linker. BRIL insertion was guided by monitoring the rigidity of the BRIL linkage using AlphaFold3, and the insertion linkage was modified using the strategy for a previously reported FZD5-BRIL fusion(40).

### Protein expression and purification of FZD4-BRIL-Tspan12 complexes

The FZD4-Tspan12 fusion protein was expressed in Expi293F TetR cells containing a stably integrated tetracycline repressor grown in Expi293 (Thermo) media supplemented with 20 μg/mL Blasticidin (Invivogen). 500 mL of Expi293F TetR cells were grown to a density of approximately 2.8 x 10^6^ cells/mL and then transiently transfected with 0.4 mg of DNA and 400 μL FectoPro transfection reagent (Polyplus). 24 hours after transfection, the cells were fed 3 mM valproic acid sodium salt (Sigma-Aldrich) and 4.5 mL of 45% D-(+)-Glucose solution (Sigma-Aldrich). 24 hours later, the transfected cells were induced with 5 mL 0.4 mg/mL doxycycline hyclate. 24 hours after induction, cells were harvested by centrifugation at 4000 x g for 10 minutes at 4°C. The mass of the pellet was recorded, and then the pellet was flash frozen and stored at −80°C until use.

The complex was purified similarly to a previously described protocol (38). Cells were thawed and then lysed by osmotic shock in 150 mL buffer containing 20 mM HEPES, pH 7.4, 2 mM magnesium chloride, 2 mg/mL iodoacetamide (Sigma Aldrich), and 1:100,000 (v:v) benzonase nuclease (Sigma Aldrich). Lysed cells were centrifuged at 18,000 × g for 20 minutes at 4°C. FZD4-Tspan12 was then extracted from the pellet using a glass dounce in 150 mL solubilization buffer containing 20 mM HEPES pH 7.4, 250 mM NaCl, 10% (v/v) glycerol, 1% (w/v) n-Dodecyl-β-D-Maltoside (DDM - Anagrade; Anatrace), 0.1% (w/v) cholesterol hemisuccinate (CHS;Steraloids), and 2 mg/mL iodoacetamide. The sample was stirred for 2 hours at 4°C and then centrifuged at 20,000 × g for 30 minutes at 4°C. The supernatant, supplemented with 2 mM calcium chloride, was filtered through a glass microfiber prefilter and loaded by gravity flow onto 2.5 mL of M1 anti-FLAG antibody affinity resin. The resin was washed with 100 mL of buffer containing 20 mM HEPES pH 7.4, 150 mM NaCl, 2 mM CaCl_2_, 0.1% DDM, and 0.01% CHS, and then 100 mL of buffer containing 2 mM CaCl_2_, 150 mM NaCl, 20 mM HEPES pH 7.4, 0.05% GDN (Anatrace), and 0.005% CHS. Bound protein was eluted in 20 mL buffer containing 20 mM HEPES pH 7.4, 150 mM NaCl, 0.05% GDN, and 0.005% CHS, 5 mM EDTA, and 0.2 mg/mL FLAG peptide. The eluate was concentrated in a 100 kDa MWCO centrifugal filter and then complexed with anti-BRIL Fab and anti-Fab nanobodies (Nb). FZD4-Tspan12 was incubated with anti-BRIL Fab and anti-Fab Nb at a 1:1.4:2 molar ratio, respectively, on ice for 1 hour. The complex was then purified by size exclusion chromatography (SEC) on a Superose 6 column (GE Healthcare) in buffer containing 20 mM HEPES pH 7.4, 150 mM NaCl, 0.05% GDN, and 0.005% CHS. Immediately after the FZD4-Tspan12-Fab-Nb complex eluted from the S6 column, two fractions were concentrated to 3.65 mg/mL using a 100k MWCO centrifugal filter and then applied to grids. Purity and monodispersity of the sample for cryo-EM were evaluated by SDS-PAGE and SEC **(Supplementary Figure 1)**.

### Purification for anti-BRIL Fab

The heavy and light sequences of the previously described anti-BRIL Fab (BAG2) (43) were cloned into the pTarget and p261040 (Atum) vectors, respectively. A set of previously described stabilizing mutations (SSASTKG replaced with FNQIKG) were introduced into the hinge region of the heavy chain to increase its rigidity (73). The Fab was expressed similarly to a previously described protocol (56). 500 mL of Expi293F cells at a density of 2.8 x 10^6^ cells/mL were transfected with 0.2 mg heavy chain DNA, 0.2 mg light chain DNA, and 200 μL FectoPro transfection reagent (Polyplus). Approximately 20 hours after transfection, cells were fed with 3 mM valproic acid sodium salt (Sigma-Aldrich) and 4.5 mL of 45% D-(+)-Glucose solution (Sigma-Aldrich) and then cultured for 5 days. The culture was harvested by centrifugation at 4000 × g for 15 minutes at 4°C. The supernatant was loaded onto 2.5 mL CaptureSelect CH1-XL Affinity Matrix resin (ThermoScientific Cat. No.1943462005). The resin was washed with 100 mL buffer containing 20 mM HEPES, pH 7.4, and 150 mM NaCl, and protein was eluted in 15 mL 100 mM citrate buffer, pH 3.0. The eluted protein was neutralized with buffer containing 1 M HEPES, pH 8. Protein was then concentrated using an Amicon 30 kDa MWCO centrifugal filter to a final volume of approximately 2 mL (17 mg/mL) and dialyzed overnight into phosphate-buffered saline (PBS). Protein purity was analyzed using an SDS-PAGE Coomassie-stained gel, and protein aliquots were flash-frozen in liquid nitrogen and stored at −80 °C. Before complexing to FZD4-Tspan12, anti-BRIL Fab and anti-Fab nanobody were buffer-exchanged into buffer containing 20 mM HEPES pH 7.4, 150 mM NaCl, 0.05% GDN, and 0.005% CHS (w/v).

### Cryo-EM sample preparation and data collection

Samples for cryo-EM were prepared on Quantifoil holey carbon film-coated 400 mesh gold grids (Electron Microscopy Sciences, R1.2/1.3, Cat: N1-C14nAu40-01). Grids were glow-discharged before the application of 3.5 μL of the sample (3.6 mg/mL). After blotting for 5.5 s with a blot force of 5, the grids were plunge-frozen in liquid ethane using a FEI Vitrobot Mark IV (FEI) with 100% chamber humidity at room temperature.

### Cryo-EM data processing and model building

Grids were loaded onto a 2^nd^-generation Titan Krios, and data collection was automated through Serial EM(74). Collection parameters are shown in Supplementary Table 1. Micrographs were imported into CryoSPARC v4.5.3(75) and processed as shown in Supplementary Figure 2A to resolve the FZD4-BRIL-Tspan12 fusion construct in complex with BRIL fab and anti-fab nanobody to 3.38 Å (best overall map) and 3.55 Å (Tspan12 focused map) using non-uniform refinement (76). The Tspan12-focused map was further processed to 3.4 Å using Deep EMhancer (77, 78).

Model building was carried out using both non-uniform refinement and deepEMhancer maps overlayed in Coot(79). The AlphaFold3 models of Tspan12 and FZD4-BRIL and the atomic coordinates of the BRIL Fab and anti-Fab nanobody (PDB: 6WW2 were manually fitted into the density map using ChimeraX to generate a starting model(80). The transmembrane domains and extracellular domains of Tspan12 docked into the density separately to allow for fitting. The coordinates were then manually rebuilt using Coot(79). TM4 of Tspan12 was poorly resolved and modeled as polyalanine, and side chains with poor density within the Tspan12 LEL or TMs with poor density were truncated to Cβ. All models were refined in Phenix RealSpace Refine (81, 82) with secondary structure restraints against the 3.4. map from non-uniform refinement in cryoSPARC. The final models were evaluated by MolProbity and EMRinger (83, 84). Statistics of the map reconstruction and model refinement are presented in Supplementary Table 1. Structural biology applications used in this project (other than CryoSPARC) were compiled and configured by SBGrid (85). Structural depictions were created through UCSF ChimeraX.

### Tspan12 trafficking assay

2 mL of Expi293F TetR cells were seeded at 2.8E6 cells/mL in a 6 well plate and then transfected using FectoPro (PolyPlus) with either: **1)** 1.5 ug empty pcDNA3.1tet/zeo (+) vector, **2)** 0.75 ug empty pcDNA3.1tet/zeo (+) vector and 0.75 ug pcDNA3.1tet/zeo (+) Tspan12, **3)** 0.75 ug pcDNA3.1tet/zeo (+) FLAG-FZD4 (full-length, WT or a chimeric construct) and 0.75 ug pcDNA3.1tet/zeo (+) Tspan12 (full-length), or **4)** 1.5 ug FLAG-FZD4-Tspan12 fusion. 24 hours after transfection, 3 mM valproic acid sodium salt (Sigma-Aldrich) and 18 μL of 45% D-(+)-Glucose solution (Sigma-Aldrich) were added to the cultures. 24 hours later, receptor expression was induced with 20 μL of 0.4 mg/mL doxycycline hyclate. 24 hours after induction, cells were harvested by centrifugation at 4000 x g for 10 minutes at 4°C. Approximately 500,000 cells were added to each well of a V-bottom 96-well plate (Costar), and cells were washed with cold PBS. Cells were then incubated on ice for 30 minutes with 10 μg/mL Alexa 488-anti-M1 FLAG (hybridoma) and Alexa 647-anti-Tspan12 (Biotechne Catalog #: FAB8910R) in 20 mM HEPES buffer pH 7.4, containing 150 mM NaCl, 2 mM CaCl2, and 0.1% BSA. Cells were washed twice with cold PBS and analyzed on an Agilent Novocyte 3000.

For western blots assessing the total cell levels of Tspan12, transfected cells were harvested by centrifugation at 4000 x g for 10 minutes at 4°C. The supernatant was discarded, and the cell pellet was lysed in 1 mL SDS loading dye supplemented with benzonase. 10 μL of lysate was run on a non-reducing SDS-PAGE gel, and the protein was transferred to a nitrocellulose membrane. The transferred protein was visualized with Ponceau staining, and then the membrane was blocked at room temperature for 1 hour in TBST (0.1% Tween-20 in Tris-buffered-saline) with 5% (w/v) Bovine Serum Albumin (BSA) (Fisher Scientific BP9706100). The blocked membrane was cut and then incubated overnight at 4°C with shaking in TBST with 1% (w/v) BSA containing either FLAG-HRP conjugate antibody (Cell Signaling 86861S; 1:1,000 dilution) or anti-Protein C (PrC) tag antibody (homemade from murine HPC4 hybridoma, 1:5000 dilution of 0.5 mg/mL stock) supplemented with 2 mM CaCl_2_. The next morning, the Tspan12-PrC blot was incubated with anti-mouse IgG HRP Conjugate (Cell Signaling 7076S; 1:5,000) in TBST with 1% (w/v) BSA for 1 hour at room temperature with shaking. Blots were washed with 10 mL of TBST three times, each for 5 minutes. Blots were developed with Western Lightning Plus-ECL, Enhanced Chemiluminescence Detection Kit (PerkinElmer), and then imaged.

### Expression and purification of MBP-Norrin

MBP-Norrin was expressed and purified as previously described (37, 59). MBP-3C-Norrin (33–133)-1D4 with an IL2 signal sequence was cloned into the pFUSE vector (Invitrogen). 500 mL of Expi293 cells were grown to a density of approximately 2.8 x 10^6^ cells/mL and then transiently transfected with 0.4 mg of DNA and FectoPro transfection reagent (Polyplus) at a 1:1 DNA/FectoPro ratio. 24 hours after transfection, the cells were fed 3 mM Valproic acid sodium salt (Sigma-Aldrich) and 4.5 mL of 45% D-(+)-Glucose solution (Sigma-Aldrich). 9 days after transfection, the culture was harvested by centrifugation at 4000x g for 15 minutes at 4°C. The culture supernatant was loaded onto 2 mL amylose resin (New England Biolabs). The resin was washed with 5 column volumes of buffer containing 20 mM HEPES pH 8, 150 mM NaCl, 1 mM EDTA, and 5% glycerol. Bound protein was eluted in 20 mL of buffer containing 20 mM HEPES pH 8, 150 mM NaCl, 1 mM EDTA, 5% glycerol, and 10 mM maltose. Eluted MBP-Norrin was concentrated using a 30 MWCO spin concentrator (Amicon), and concentrated protein was loaded on a Superdex 200 Increase 10/300 GL equilibrated in buffer containing 20 mM HEPES pH 8, 150 mM NaCl, 1 mM EDTA, and 5% glycerol. Peak fractions were pooled, concentrated to approximately 1 mg/mL, flash frozen, and frozen in 25 μL aliquots at −80°C until use **(Supplementary Figure 12)**. MBP-Norrin was validated in FZD4 binding assays before use.

### MBP-Norrin binding assays

2 mL of Expi293F TetR cells were seeded at 2.8E^6^ cells/mL in a 6 well plate and then transfected using FectoPro (PolyPlus) with either 1.5 μg empty pcDNA3.1tet/zeo (+) or 1.5 ug of FLAG-apo-FZD4, FLAG-FZD4-BRIL-Tspan12, E170K FLAG-FZD4-BRIL-Tspan12, or FLAG-FZD4 with the CRD and hinge of FZD10. 24 hours after transfection, cells were fed with 3 mM valproic acid sodium salt (Sigma-Aldrich) and 18 μL of 45% D-(+)-Glucose solution (Sigma-Aldrich). 24 hours later, expression was induced with 20 μL of 0.4 mg/mL doxycycline hyclate (dissolved in buffer containing 20 mM HEPES pH 7.4, 150 mM NaCl). 24 hours later, cells were harvested by centrifugation at 4,000 x g at 4 °C and resuspended in cold phosphate buffered saline (PBS). The cells were then counted, and 100,000 cells were added to each well of a V-bottom 96-well plate (Grenier Bio). Cells were washed again with 100 μL cold PBS and then incubated with different concentrations of either the 1D4-tagged MBP-Norrin, secondary antibody alone (anti-1D4-488),

M2-FLAG-488 to quantify receptor expression level, or buffer alone for thirty minutes at 4°C. Following another PBS wash, cells were incubated with 2 μg/mL anti-Rho1D4-Alexa Fluor 488 antibody for 30 minutes at 4°C to detect Norrin binding. After secondary antibody incubation, cells were washed twice more with PBS and were analyzed by flow cytometry using a Novocyte 3000 (Agilent). Live/dead gates were set using SSC-A versus FSC-A and singlet gates using FSC-H versus FSC-A, and fluorescence intensity in the 488 channel was recorded in the doubly gated population. Median fluorescence intensity (MFI) at 488 nm was plotted in GraphPad Prism as an assessment of MBP-Norrin binding to cells. For EC50 calculations, non-specific signal was subtracted (MFI in the mock condition treated with the corresponding MBP-Norrin concentration). Curve fitting was performed in Prism (GraphPad, v10) to a log (agonist) vs. response four-parameter model, with no constraints on Hill Slope, EC50, or top values, and a constraint of 0 on the bottom value.

### BRET assays

ΔFZD_1–10_ HEK293T cells were used for all BRET experiments. White 96-well tissue-culture treated plates (Revity) were precoated with 50 μL/well of a 0.1 mg/mL poly-D-lysine (Sigma) and incubated at room temperature for 20 minutes. The coating solution was aspirated and washed once with 200 μL/well of DPBS, and the plate was dried for 20 minutes in the tissue culture hood before seeding cells. ΔFZD_1–10_ HEK293T cells were trypsinized and diluted to 0.4E^5^ cells/mL. 1 mL of cells at 0.4E^6^/mL were then transfected in suspension. Acceptor: donor transfection ratios were pre-determined from acceptor: donor titration experiments to determine the signal plateau and fold increase over the CD151-Venus negative control. For transfections, Tube A contained 40 μL opti-MEM, 20 ng FZD4-NLuc, 100 ng Tspan12-Venus or CD151 Venus, or 0 ng of a Venus construct as a background control, and 20 ng untagged LRP5. An empty pcDNA3.1 vector was used to bring the final DNA concentration to 1,000 ng. Tube B contained 49 μL Opti-MEM and 1 μL Lipofectamine 2000. Tubes A and B were incubated separately for 5 minutes and then added together and incubated at room temperature for 10 minutes. The transfection mix was added dropwise to 1 mL of cells at 0.4E^6^/mL, 100 μL of the cell-transfection mix was added to each well of the poly-D-lysine-coated 96-well plate. Approximately 24 hours after transfection, the cells were washed once with 200 μL of HBSS and then kept in HBSS. 10 μL of furimazine (Promega) (final dilution at 1:1000) was added to a final volume of 90 μL, and 5 min after furimazine addition, luminescence was read three times for a baseline reading. Subsequently, 10 μL of Norrin at a final concentration of 10 nM (in HBSS) was added to a final volume of 100 μL, and luminescence reading was continued 44 times for a total of 54.8 minutes. All measurements were performed at 37°C using a ClarioStar plate reader. BRET experiments were performed with five replicates.

### Tspan12 Pulldown

ΔFZD_1–10_ HEK293T cells were seeded at 0.3E^6^ cells/mL in 10 mL DMEM with 10% FBS (Fisher Scientific SH3007003HI) and 1% Pen-Strep (Thermo Scientific 15140122). Approximately 12 hours later, media was replaced with 10 mL DMEM with 2% FBS and no Pen/Strep. Per 10 cm^3^ plate, cells were transfected with 1.2 ug DNA, at a ratio of 1:1.5:4 FZD4:LRP5:Tspan12, or 1.2 μg empty vector (pcDNA3.1), and 40 μL lipofectamine 2000 in 1.5 mL opti-MEM. The FZD4 construct was N-terminally FLAG tagged, Tspan12 was N-terminally Protein C (PrC) tagged, and LRP5 was untagged. The transfection mix was incubated for 15 minutes at room temperature and then added dropwise to the cells. The next day, the media was replaced with DMEM with 10% FBS and 1% Pen-Strep. 48 hours after transfection, cells were serum starved for 1 hour in DMEM with no FBS and then treated with 10 nM of Norrin for 15 minutes or left untreated. Cells were harvested in 2 mL cold PBS with 5 mM EDTA, pelleted at 4000 x g for 10 minutes at 4°C.

Pulldowns were performed as previously described (56). Briefly, the pellet was lysed by osmotic shock in buffer containing 20 mM HEPES pH 7.4, 2 mM MgCl_2_, 2 mg/mL iodoacetamide, and 1:100,000 v:v benzonase. Samples were rotated for 30 minutes at 4°C and then centrifuged at 16,000 × g for 15 minutes at 4°C. Solubilization buffer (1% (w/v) DDM/0.1% CHS buffer, 20 mM HEPES pH 7.4, 250 mM NaCl, 10% glycerol, 2 mg/mL iodoacetamide) was added to the pellet, and then the samples were rotated for 2 hours at 4°C. Samples were pelleted at 20,000 x g for 30 minutes at 4°C and then added to 25 μL of M2 magnetic FLAG beads (Sigma, M8823). The samples were rotated for 2 hours in the cold room, and then beads were washed three times with 1 mL of buffer containing 0.1% (w/v) DDM, 0.01% CHS buffer, 20 mM HEPES pH 7.4, 150 mM NaCl. Proteins bound to the beads were eluted in 250 μL SDS loading dye supplemented with 0.2 mg/mL FLAG peptide.

Western blot analysis was performed similarly to a previously described protocol (56). 10% input and 10% eluate samples were run on a non-reducing SDS-PAGE gel, and then the protein was transferred to a nitrocellulose membrane. Next, the membrane was blocked in TBST with 5% (w/v) BSA (Fisher Scientific BP9706100) at room temperature for 1 hour, shaking. The membrane was then cut and incubated in TBST with 1% (w/v) BSA containing either FLAG-HRP conjugate antibody (Cell Signaling 86861S; 1:1,000 dilution) or anti-Protein C (PrC) tag antibody (Cell Signaling 68083; 1:1000 dilution), overnight at 4°C with shaking. The next morning, the Tspan12-PrC blot was incubated with anti-Rabbit IgG HRP Conjugate (Cell Signaling 7074S; 1:5,000) in TBST with 1% (w/v) BSA for 1 hour at room temperature with shaking. Blots were washed three times with 10 mL of TBST and then developed with Western Lightning Plus-ECL, Enhanced Chemiluminescence Detection Kit (PerkinElmer) prior to imaging.

### TOPFlash Signaling Assays

ΔFZD1–10 HEK293T were used for all TOPFlash assays. White 96-well tissue-culture treated plates (Revity) were precoated with 80 μL/well of a 0.1 mg/mL poly-D-lysine solution (Sigma) and incubated at room temperature for 20 minutes. The coating solution was aspirated and washed once with 100 μL/well of DPBS. The plate was then dried for 20 minutes in the tissue culture hood before seeding the cells. For tethered FLAG-FZD4-Tspan12 versus FLAG-FZD4 and Tspan12 co-transfections (Figure 4C), Tube A contained 250 uL of OptiMEM, 250 ng of Super 8x TOPFlash plasmid M50 (Addgene #12456), 50 ng Renilla luciferase control plasmid (pRL-TK, Promega), 250 ng of LRP5, 10 μg of FLAG-FZD4-Tspan12 or co-transfections of 200 ng FLAG-FZD4 with 600 ng Tspan12. For all other constructs tested, Tube A contained 250 uL of OptiMEM, 250 ng of Super 8x TOPFlash plasmid M50 (Addgene #12456), 50 ng Renilla luciferase control plasmid (pRL-TK, Promega), 50 ng of LRP5, 50 ng FLAG-FZD4 with 250 ng Tspan12. Empty pcDNA3.1 vector was used to bring to a final concentration of DNA to 2,500 ng for Tspan12 and FZD4 co-transfections. Empty pcDNA3.1 vector was used as a control and included Super 8x TOPFLASH plasmid M50 and Renilla luciferase control plasmid (pRL-TK). Tube B contained 250 μL of OptiMEM with Lipofectamine 2000 (2 μL per 1 μg of DNA). Tubes A and B were incubated separately for 5 minutes and then combined to incubated for 15-30 minutes. ΔFZD1–10 HEK293T cells were trypsinized and diluted to 0.5E^6^ cells/mL with DMEM (10% FBS, 1% PenStrep). 2.5 mL of cells at 0.5E^6^/mL were then added to the combined Tube A+B incubation. 100 μL of transfected cells were added to the poly-D-lysine coated plates (12 wells per construct) and 1.2 mL of transfected cells were plated in 12-well plates for companion flow staining assay (described below). 24 hours post-transfection, media was aspirated and replaced with DMEM (0% FBS, no PenStrep) with 10 nM C59-porcupine inhibitor (VWR) that contained either 0 nM MBP-Norrin (control), 1 nM MBP-Norrin, 10 nM MBP-Norrin, or 25 nM MBP-Norrin, in triplicate. For the companion flow assay to detect receptor expression levels, the media was swapped with DMEM (0% FBS, no PenStrep) with 10 nM C59-porcupine inhibitor without Norrin. 24 hours after media swap, media was aspirated and cells were washed with 100 μL DPBS (Corning). The Dual

Luciferase Assay Kit (Promega, #E1910) was used for TopFlash readout. Cells were lysed with 20 μL of 1x Passive Lysis Buffer for 20 minutes while shaking at room temperature. A ClarioStar plate reader with dual injector was used to inject 20 μL of LARII reagent into each well and firefly luciferase bioluminescence was recorded (580 ± 80 nm range and 10 second interval time). This was immediately followed by injection of 20 μL of 1x Stop-and-Glo reagent per well and Renilla luciferase bioluminescence was recorded (480 ± 80 nm range and 10 second interval time).

For FZD4/Tspan12 constructs used in the TOPFlash assay, expression levels were monitored using flow cytometry. 36-48 hours after transfection (described above), and 18-24 hours after DMEM (0% FBS, no PenStrep) media swap, the media was aspirated from the 12-well plates, and 1.0 mL of cold PBS + 3 mM EDTA was added to the wells and mixed well to suspend the cells. 200 μL of cells were added to 4 wells in 96-well v-bottom clear flow plates, kept on ice. Cells were spun down at 300 g for 3 minutes and then washed with 100 μL of cold PBS, followed by a second spin down. Cells were then suspended and double-stained with anti-FLAG M2 antibody conjugated to Alexa Fluor 488 (final concentration, 1 μg/μL) and anti-Tspan12 antibody conjugated to Alexa Fluor 647 (final concentration, 1 μg/μL) in HBS + 0.2% BSA for 30 minutes on ice, protected from light. Unstained control samples were kept in HBS + 0.2% BSA and no antibodies were included. After incubation, cells were washed spun down and washed 2x with cold PBS before final suspension in cold PBS+3mM EDTA+1% BSA. Fluorescence was analyzed on an Agilent Novocyte 3000.

### Flow Cytometry on Fixed and Permeabilized Cells

ΔFZD1–10 HEK293T cells were seeded at 0.3E6 cells/mL in 2 mL DMEM with 10% FBS (Fisher Scientific SH3007003HI) and 1% Pen-Strep (Thermo Scientific 15140122) in 6-well plates (CytoOne). Approximately 12 hours later, media was replaced with 2 mL DMEM with 2% FBS and no Pen/Strep. Cells were transfected with 0.3 μg DNA, at a ratio of 1:1.5:4 FZD4:LRP5:Tspan12, or 0.3 μg empty vector (EV) (pcDNA3.1), and 10 μL lipofectamine 2000 in 500 μL opti-MEM. 12 hours after transfection, media was replaced with 2 mL DMEM with 10% FBS. 48 hours after transfection, cells were harvested with 1 mL cold PBS +5 mM EDTA and placed in 1.5 mL Eppendorf tubes. Cells were spun down at 300g, for 3 min (4°C) to pellet the cells. Media was aspirated, and the cell pellet was resuspended in 1.3 mL of cold PBS. Cells were then divided into two 96-well V-bottom plates (Costar) – one for cell surface staining and the other for permeabilization and whole cell staining.

For surface staining, 100 µL of EV or Tspan12+FZD4+LRP5 transfected cells were plated into 6 wells of a 96-well v-bottom plates (Costar) and spun down at 300g, for 3 min, (4°C) washed with 150 µL cold PBS. In triplicate, EV or Tspan12+FZD4+LRP5 were double-stained with an anti-M2-FLAG antibody conjugated to Alexa Fluor-488 (2 µg/mL) and an anti-Tspan12 antibody conjugated to Alexa Fluor-647 for 30 minutes on ice and protected from light. Control samples, also in triplicate, were incubated with HBS + 2% BSA under the same conditions. After incubation, cells were spun down and washed twice with PBS before final suspension in 100 µL of cold PBS. For permeabilized cell staining, 100 µL of EV or Tspan12+FZD4+LRP5 transfected cells were plated into 6 wells on a 96-well v-bottom plates (Costar) and then spun down at 300g, for 3 min (4°C) and washed with 150 µL cold PBS. We then resuspended and fixed the cells in 100 µL PBS + 1 mM CaCl_2_ + 0.01% Formaldehyde while slowly shaking for 15 minutes at RT. This was followed by two rounds of 100 µL PBS washes and then permeabilization with 100 µL of PBS + 0.5% Tween-20 while slowly shaking for 15 minutes at RT. After two PBS washes, EV or Tspan12+FZD4+LRP5 were double-stained with an anti-M2-FLAG antibody conjugated to Alexa Fluor-488 (2 µg/mL) and an anti-Tspan12 antibody conjugated to Alexa Fluor-647 for 30 minutes on ice and protected from light, in triplicate. Control samples, also in triplicate, were incubated with HBS + 2% BSA under the same conditions. After incubation, the cells were spun down and washed twice with PBS before being resuspended in 100 µL of cold PBS. Fluorescence was analyzed on an Agilent Novocyte 3000. Live/dead gates were set using SSC-A versus FSC-A and singlet gates using FSC-H versus FSC-A, and fluorescence intensity in the 488 and 647 channels was recorded in the doubly gated population.

