## Supplementary Information for "Structural basis for regulation of Frizzled-4 signaling by the co-receptor Tetraspanin-12"

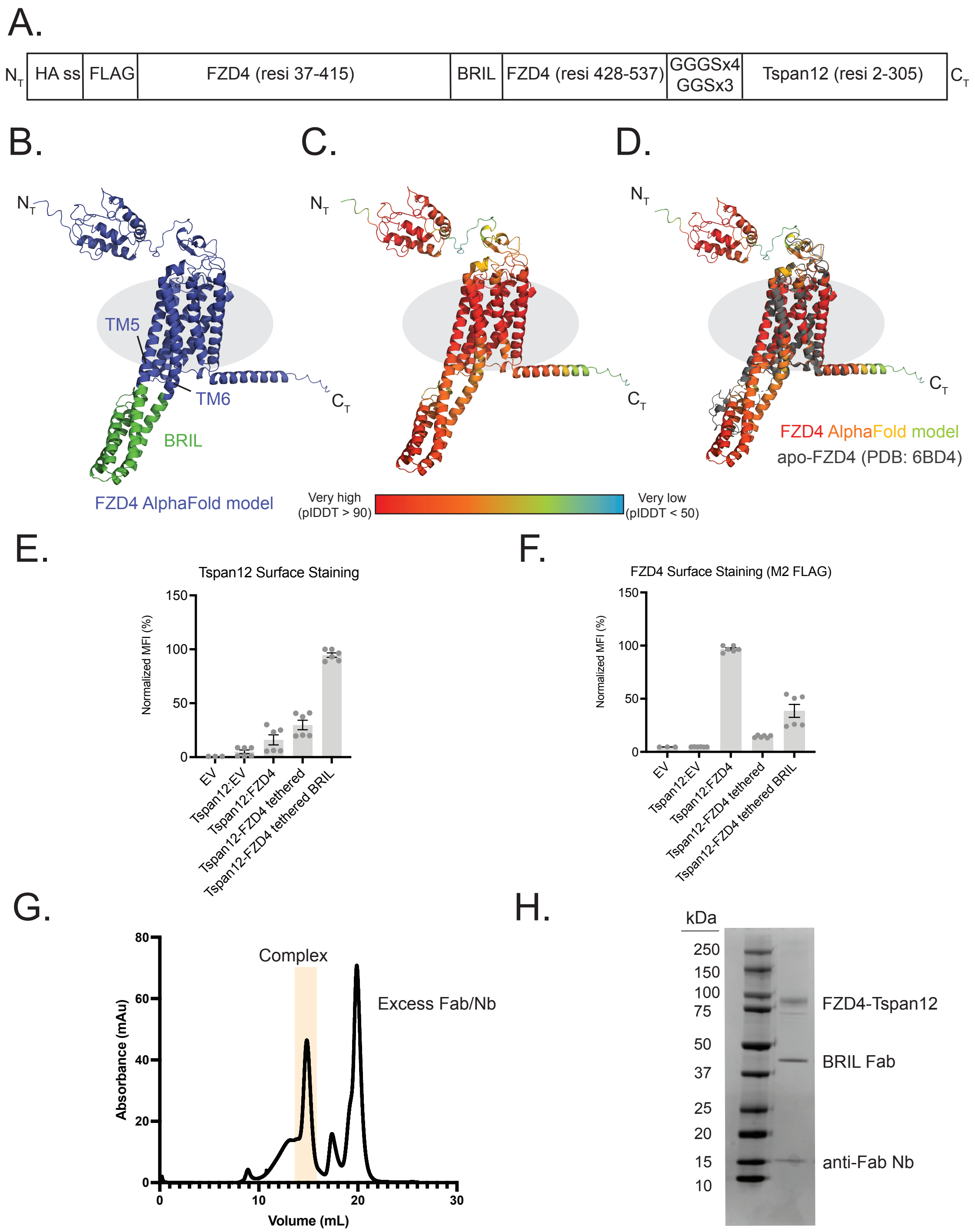


**Supplementary Figure 1: Design and validation of FZD4-BRIL-Tspan12 fusion protein. (A)** Schematic of the construct used to produce the FZD4-Tsapn12 complex. A BRIL domain was inserted between residues 415 and 428 of FZD4 and was linked to full length, wild type Tspan12 using a Gly-Gly-Ser repeat. **(B)** AlphaFold3 model of FZD4 with the BRIL domain, with the BRIL domain colored in green. **(C)** AlphaFold3 model of FZD4 with the BRIL domain colored by pIDDT score, highlighting the confident prediction of rigid helical extensions between the BRIL domain and FZD4. **(D)** Overlay of the AlphaFold3 model of FZD4 with the BRIL domain, with the crystal structure of inactive state FZD4 (PDB: 6BD4, colored in grey), showing that the BRIL domain did not result in structural changes in FZD4. **(E)** Flow cytometry-based Tspan12 trafficking assay with the FZD4-BRIL-Tspan12 fusion, showing increased surface expression of Tspan12, detected with an anti-Tspan12 antibody (BioTechne Cat #FAB8910R-025). Error bars represent mean (±) SEM of two independent experiments with three technical replicates each. **(F)** Flow cytometry-based Tspan12 trafficking assay with the FZD4-BRIL-Tspan12 fusion, monitoring FLAG-FZD4 surface expression with an anti-FLAG M2 antibody. Because FZD4 is tethered to Tspan12, Tspan12 limits surface trafficking of apo-FZD4. Error bars represent mean (±) SEM of two independent experiments with three technical replicates each. **(G)** Size exclusion profile of the purified FZD4-Tspan12-BRIL Fab-Anti-Fab nanobody complex in detergent buffer on a Superose 6 column. The peak highlighted in yellow was used for grid freezing. **(H)** SDS-PAGE gel of the purified complex.

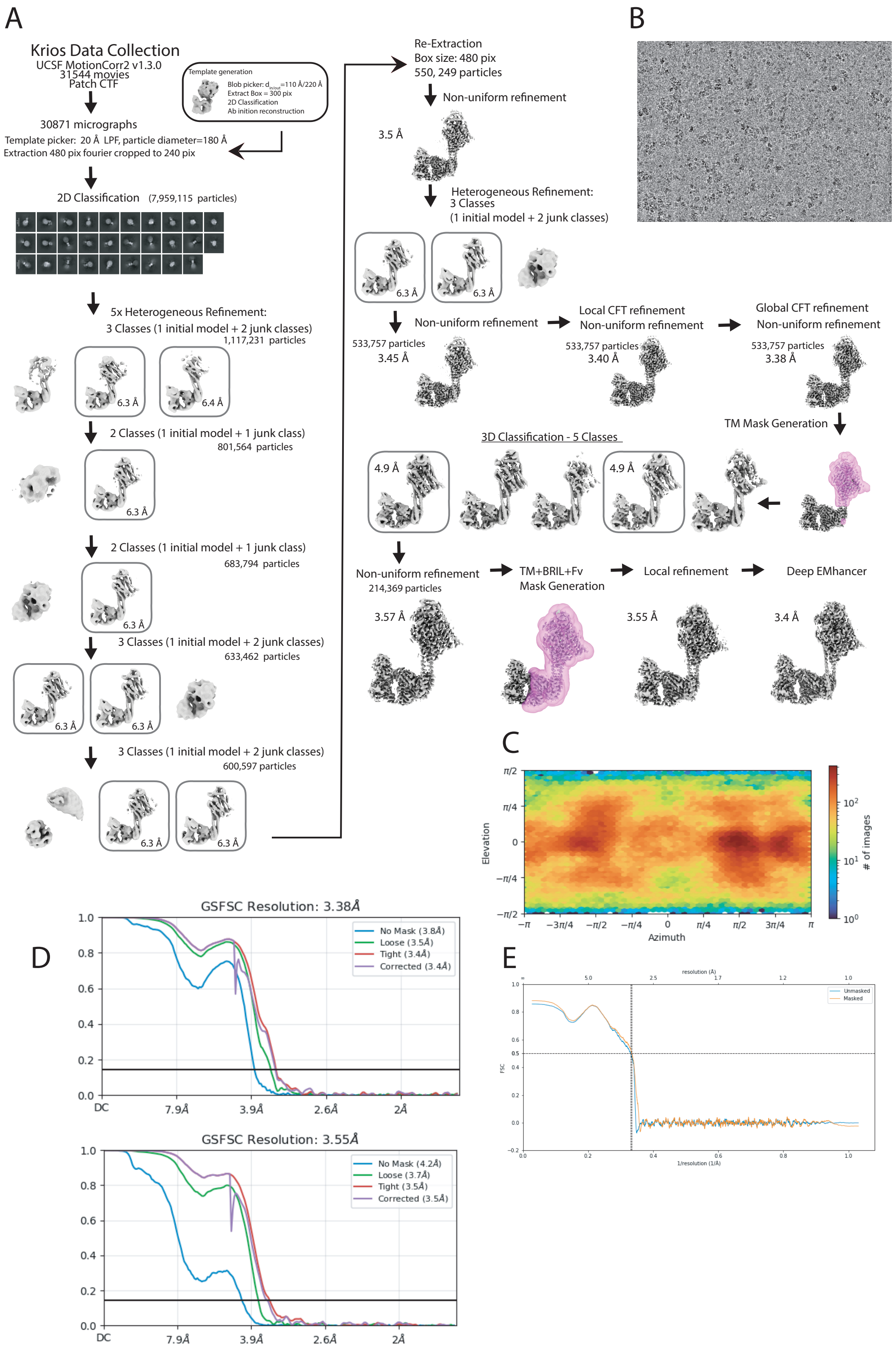


**Supplementary Figure 2:** Cryo-EM image processing workflow. **(A)** Image and particle processing steps as conducted in CryoSPARCv5.4.3 for Tspan12-Fzd4-BRIL in complex with anti-

BRIL antibody and anti-fab nanobody. **(B)** Representative cryo-EM micrograph. **(C)** Angular distribution of Tspan12-Fzd4-BRIL-Fab-Nanobody particles in the final round of non-uniform refinement in CryoSPARC. **(D)** Fourier shell correlation (FSC) of the final map (3.4 Å), and the focused Tspan12 map (3.5 Å). The black line marks the resolution corresponding to an FSC value of 0.143. **(E)** Map to model FSC curve.

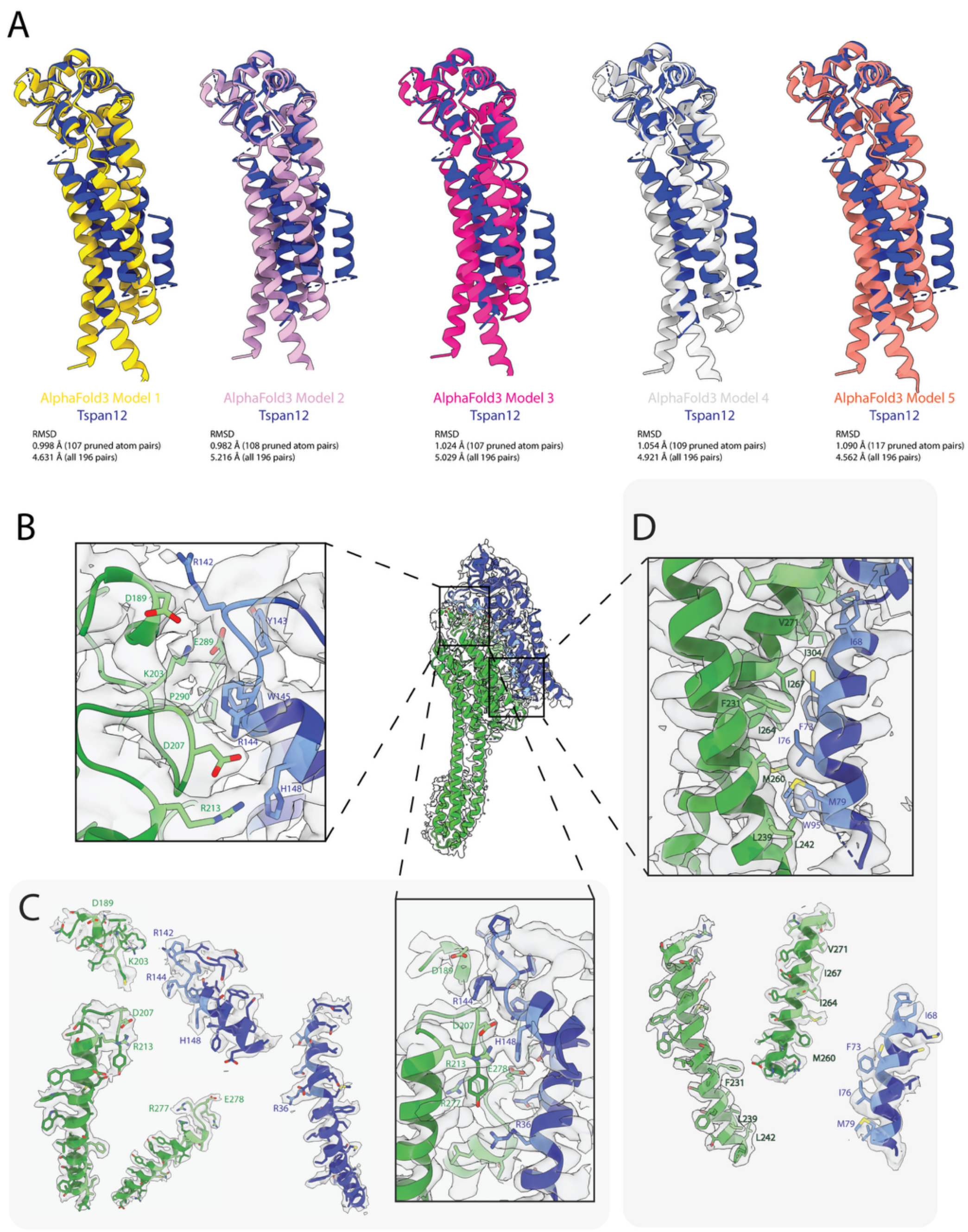


**Supplementary Figure 3:** Fit of the atomic model into the cryo-EM map. The model is shown as sticks in its corresponding cryo-EM density. **(A)** Alphafold3 predicted Tspan12 models (1-5) overlayed on Tspan12-Fzd4-BRIL structure and corresponding RMSD values. **(B)** Density overlap of electrostatic interactions between Fzd4 lower linker and the resolved LEL of Tspan12. On the right, the overall representative figure of the Tspan12-FZD4 model and map overlay is shown. Interacting residues are shown in light green (Frizzled4) and light blue (Tspan12). **(C)** Electrostatic interactions between Fzd4 lower linker, TM1 and TM2 and the EC loops of Tspan12 (SEL and LEL). Segmented view of side chain densities are depicted on the right side of the panel. **(D)** Density overlap of the hydrophobic pocket interactions between Fzd4 TM2 and Tspan12 TM2 hydrophobic residues as well as segmented view of side chain density.

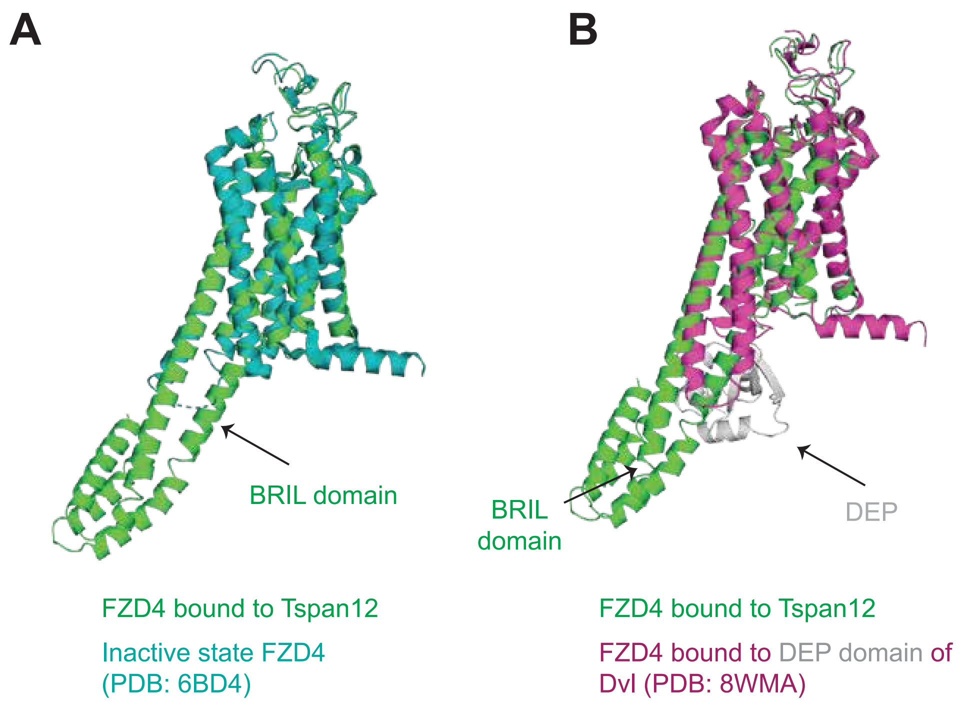


**Supplementary Figure 4:** Comparison of Tspan12-bound FZD4 to inactive state FZD4 (PDB: 6BD4) and DEP-bound FZD4 (PDB: 8WMA). Ribbon diagrams showing superposition of the structure of Tspan12-FZD4-BRIL (green) on the structure of inactive state FZD4 **(A)** (teal, RMSD: 0.682 Å) and DEP-bound FZD4 **(B)** (magenta and grey, RMSD: 1.15 Å)

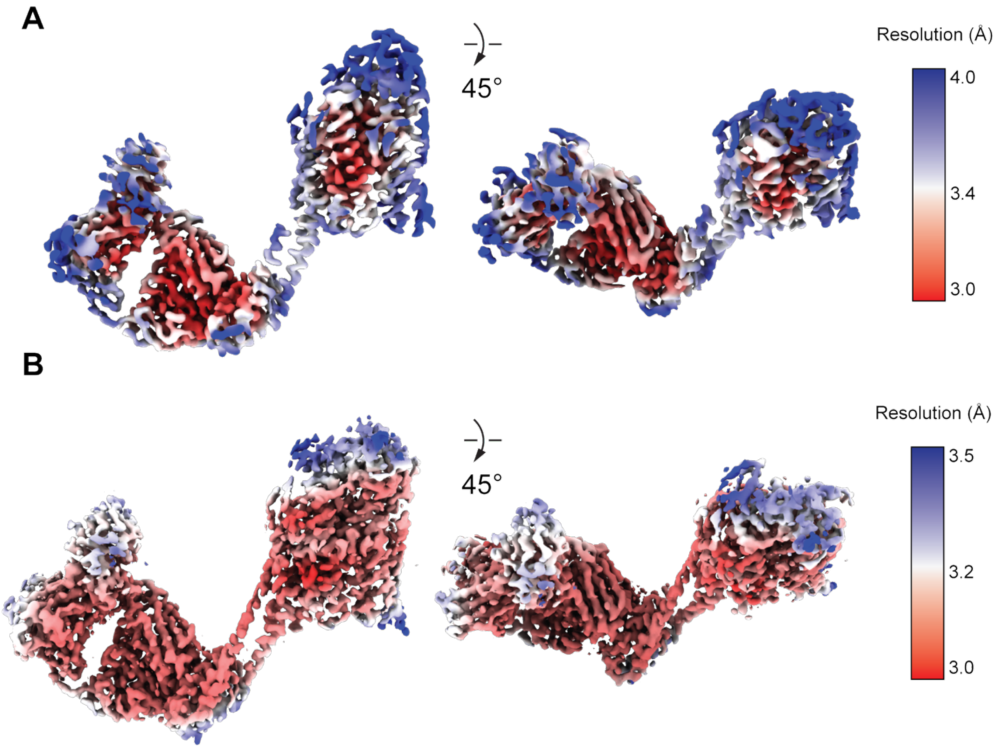


**Supplementary Figure 5: (A)** Locally filtered cryo-EM map of the Tspan12-focused map of the Tspan12-FZD4-BRIL complex colored according to local resolution, calculated in cryoSPARC. **(B)** Tspan12-focused map of the Tspan12-FZD4-BRIL complex after post-processing in deepEMhancer colored by local resolution, calculated in Phenix, resulting in a map with improved interpretability and clearer features.

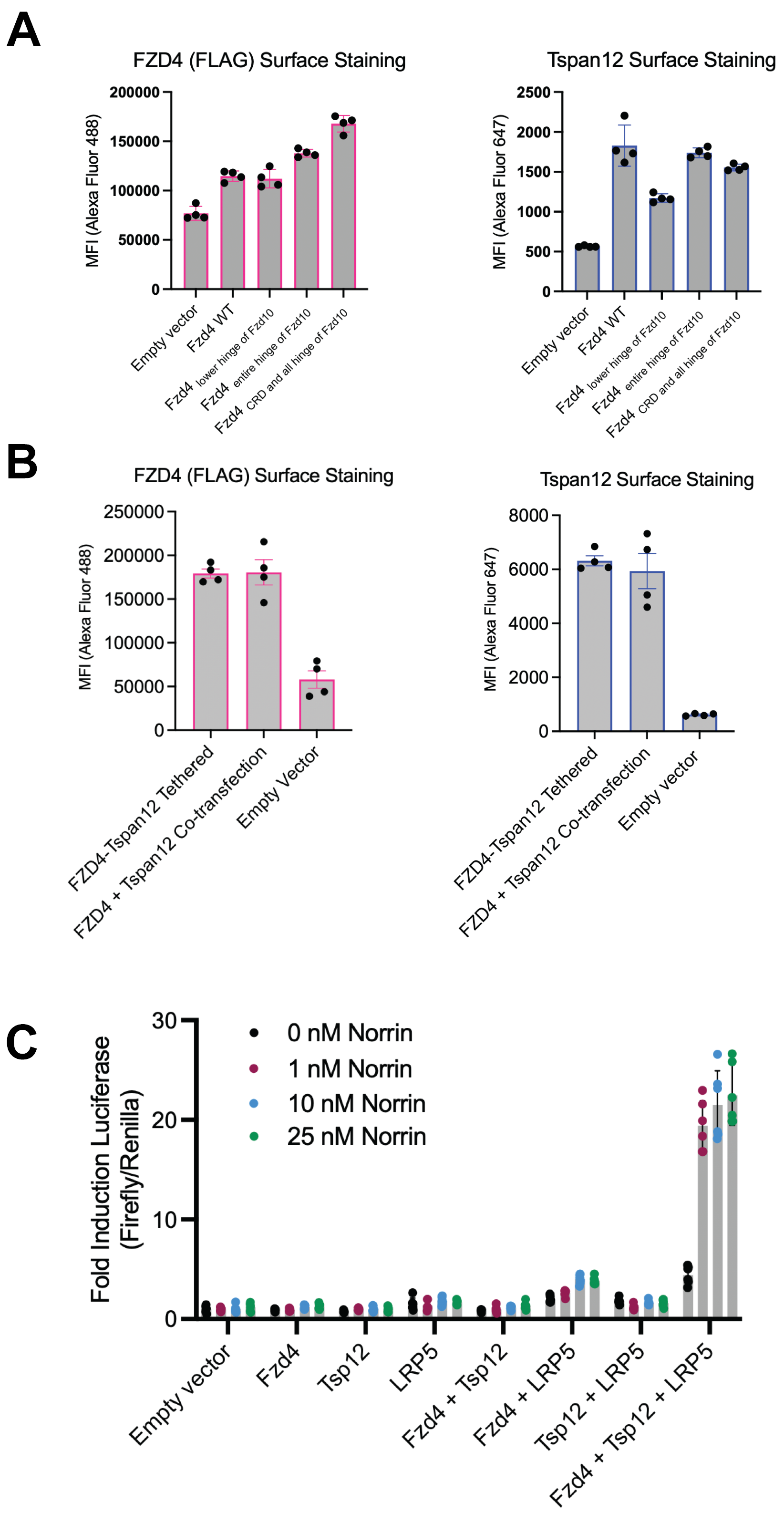


**Supplementary Figure 6: TOPFlash expression and assay controls. (A)** M2-Flag (left-panel, in pink) and Tspan12 (right panel, in blue) staining of $\Delta FZD_{1-10}$ HEK293T transfected under TOPFlash conditions with co-transfections of either FZD4-WT or FZD4-FZD10 chimeras with

Tspan12, LRP5, Super 8x TOPFLASH M50, and Renilla luciferase (pRL-TK) plasmids. Error bars represent mean (±) SEM of two independent experiments with two technical replicates each. **(B)** M2-Flag (left-panel, in pink) and Tspan12 (right panel, in blue) staining of $\Delta FZD_{1-10}$ HEK293T transfected under TOPFlash conditions comparing co-transfected FZD4-WT and Tspan12 or tethered FZD4-Tspan12 in the presence of LRP5, Super 8x TOPFLASH M50, and Renilla luciferase (pRL-TK) plasmids. Error bars represent mean (±) SEM of two independent experiments with two technical replicates each. **(C)** TOPFlash assay comparing signal with single, double and triple receptor transfections in the presence of 0 nM, 1 nM, 10 nM or 25 nM MBP-Norrin. Error bars represent mean (±) SEM of two independent experiments, one with two technical replicates and the other with three technical replicates.

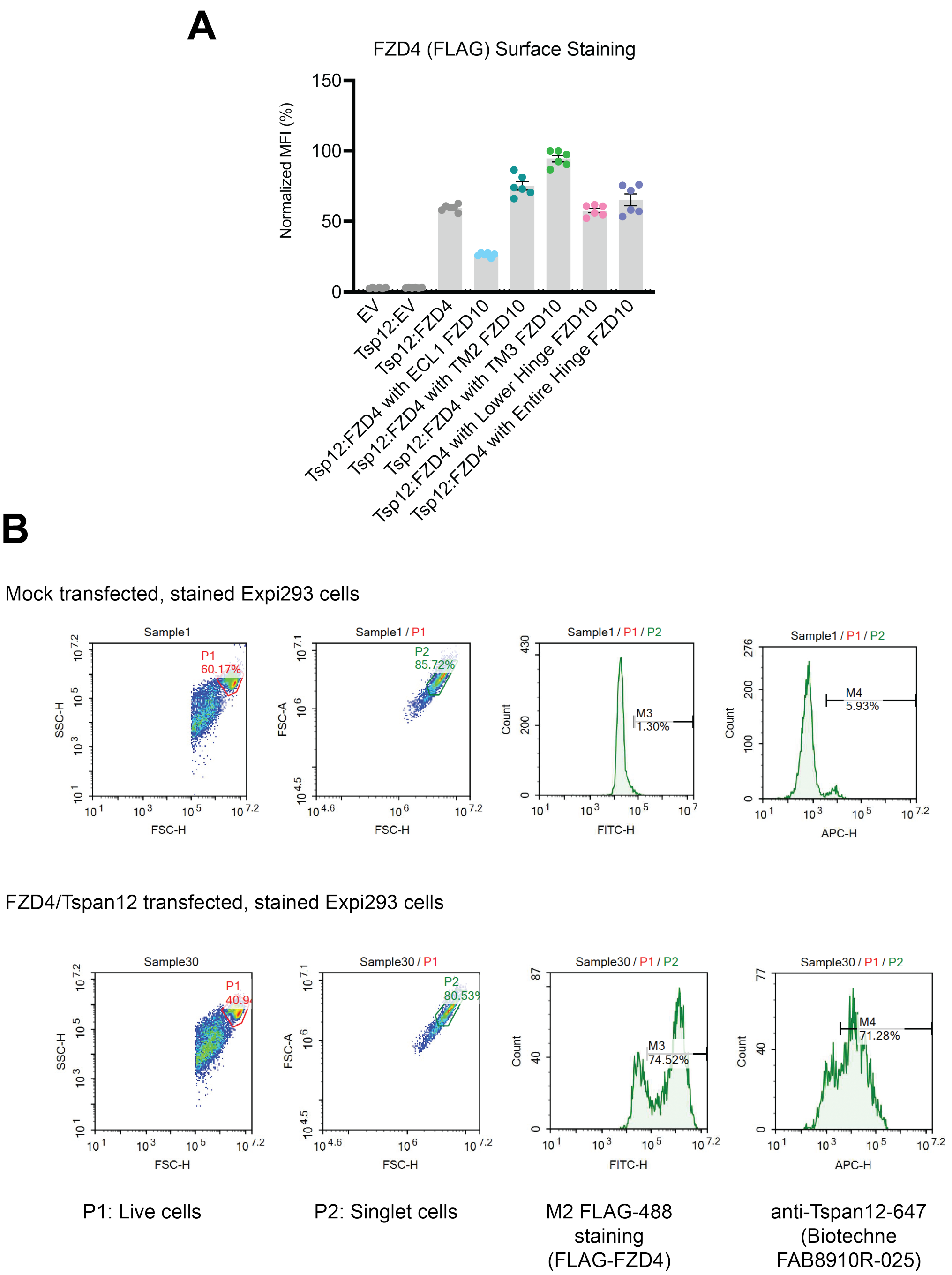


**Supplementary Figure 7:** Expression of FZD4 chimeras and representative flow cytometry gating. Error bars represent mean (±) SEM of two independent experiments with three technical replicates each. **(A)** Surface expression of FLAG-tagged FZD4 chimeras used in the Tspan12 trafficking assay, detected with an anti-FLAG M2-Alexa Fluor 488 antibody. **(B)** Representative gating strategy on Expi293 cells.

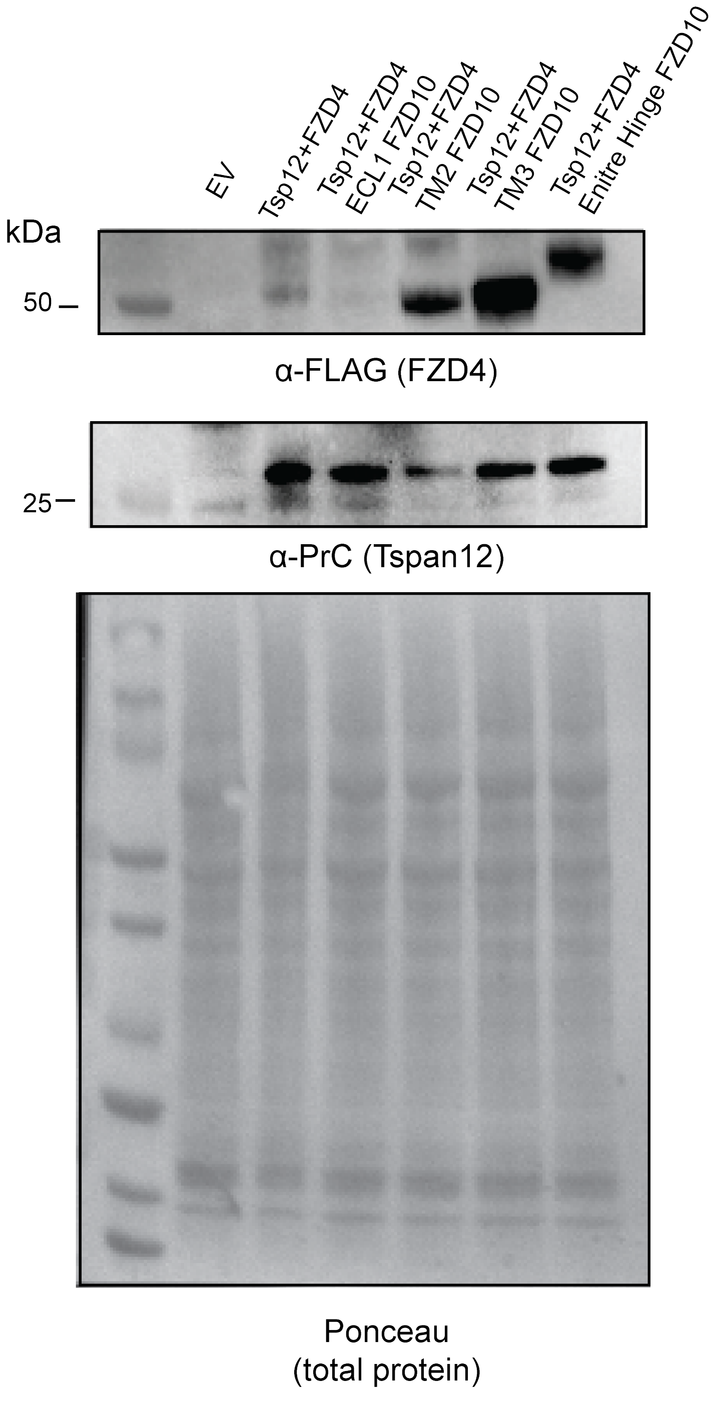


**Supplementary Figure 8: (A)** anti-FLAG (FZD4) and anti-Protein C (Tspan12) Western blots on whole cell lysates of cells transfected with Tspan12 and wild type FZD4 or the chimeric FZD4 constructs.

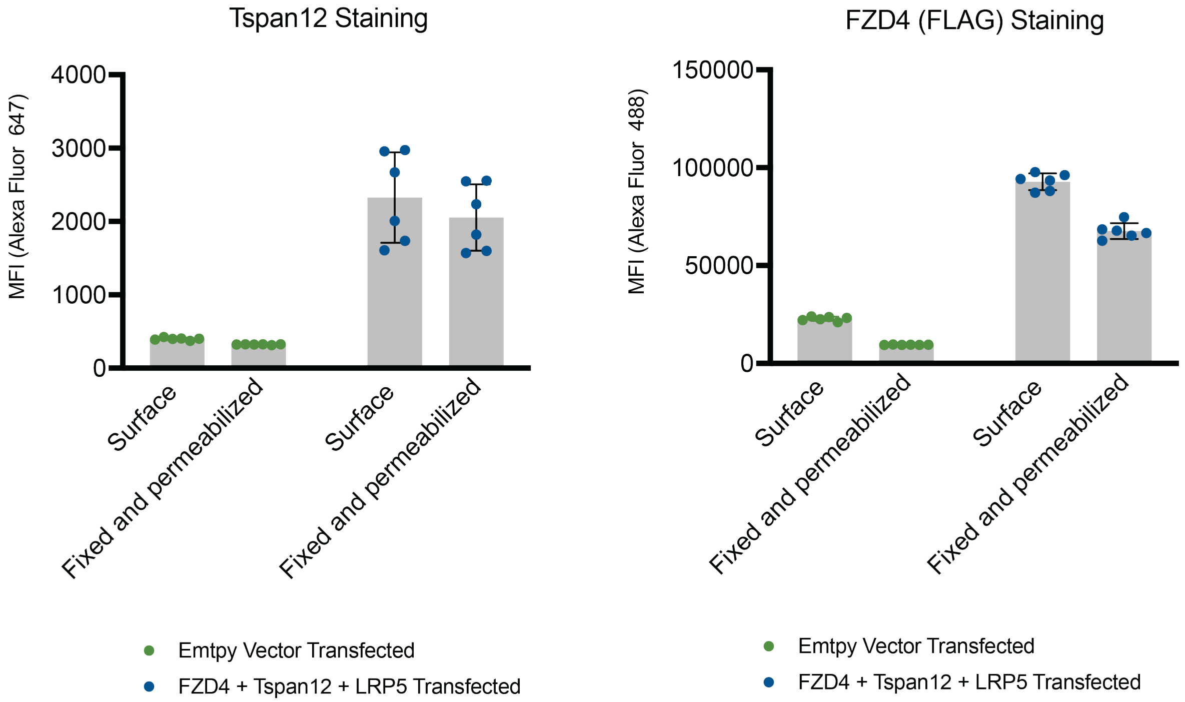


**Supplementary Figure 9:** Surface and fixed and permeabilized flow cytometry to detect relative surface and intracellular expression levels of transfected FZD4 and Tspan12 in $\Delta FZD_{1-10}$ HEK293T cells used for pulldown assays. Error bars represent mean (±) SEM of two independent experiments with three technical replicates each.

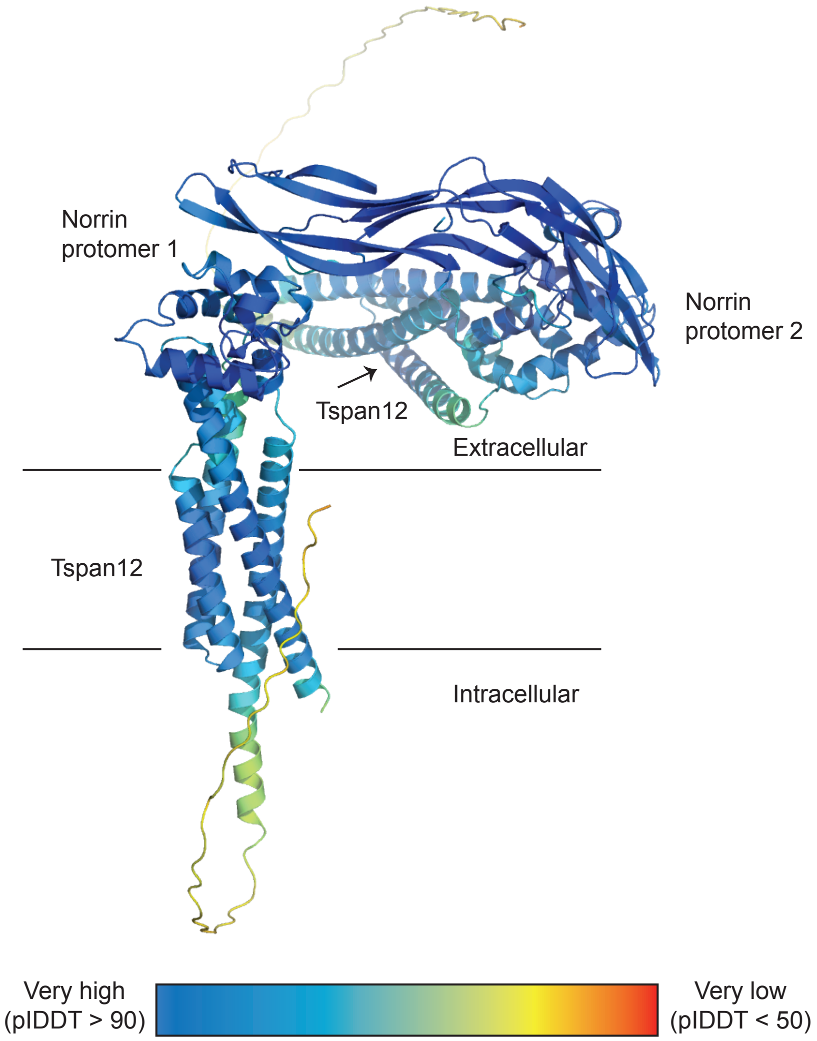


**Supplementary Figure 10:** Norrin-Tspan12 AlphaFold3 Multimer model with Tspan12 bound to each protomer of Norrin. The model is colored by pIDDT score.

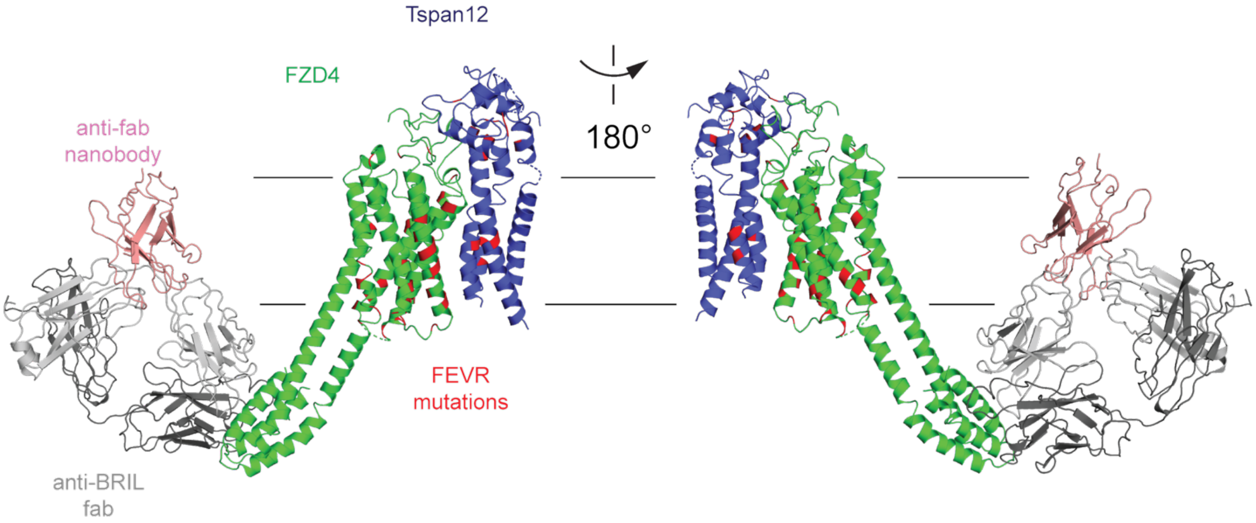


**Supplementary Figure 11:** Mapping of known FEVR missense mutations onto the structure of Tspan12-FZD4. Mutations are colored in red, Tspan12 in blue, and FZD4 in green.

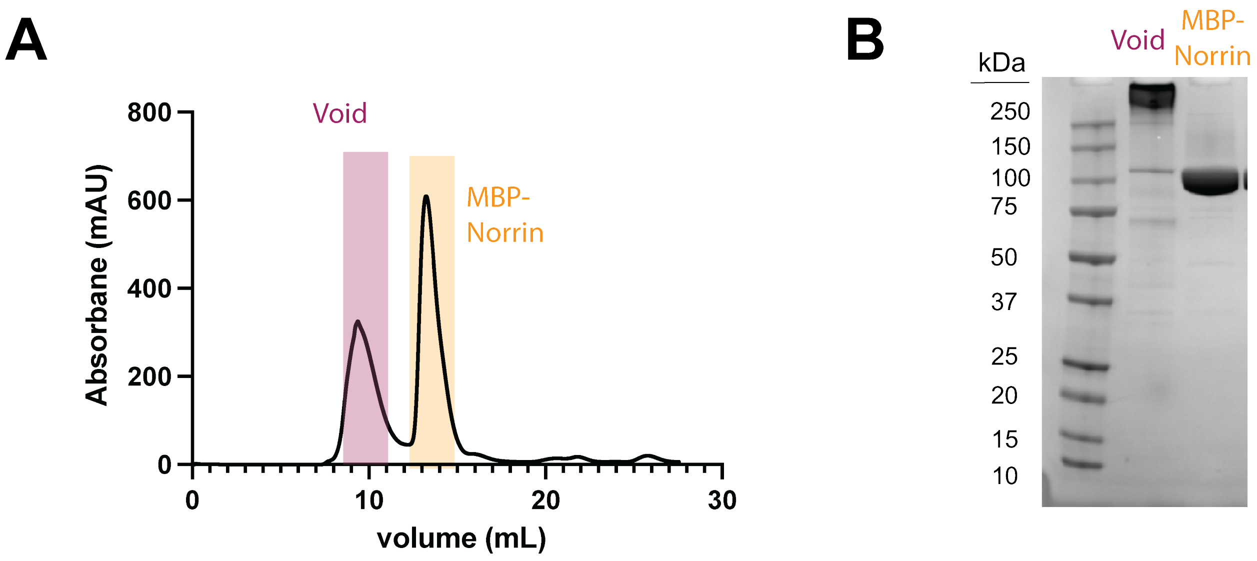


**Supplementary Figure 12:** Purification of MBP-Norrin. **(A)** Size exclusion profile of the purified MBP-Norrin on a S200 Increase column. The peak highlighted in yellow was used for binding assays and cell-based experiments. **(B)** SDS-PAGE gel of purified MBP-Norrin.

**Supplementary Table 1:** Cryo-EM data collection, refinement, and validation statistics

| | **Tspan12-FZD4 complex with anti-BRIL Fab-anti-Fab nanobody**<br>PDB: 9YEV<br>EMDB: EMD-72864 |
|---|---|
| **Data collection and processing** | |
| Microscope | Titan Krios |
| Magnification | 105,000x |
| Voltage (kV) | 300 |
| Camera | GATAN K3 |
| Pixel size at detector (Å) | 0.8189 |
| Total electron exposure (e–/ Å) | 47.7 |
| Exposure rate (e-/pixel/sec) | 16 |
| Defocus range (μm) | -1.2 to -2.2 |
| Automation software | Serial EM |
| Symmetry | C1 |
| Micrographs collected (no.) | 31, 544 |
| Micrographs used (no.) | 30, 871 |
| Total extracted particle images (no.) | 214, 369 |
| Map resolution (Å)<br>FSC threshold | 3.4<br>(0.143) |
| Map sharpening B factor ($Å^2$) | -122.2 |
| **Refinement** | |
| Initial model | AlphaFold3 model of Tspan12, AlphaFold3 model of FZD4-BRIL, and PDB 6WMA |
| Refinement package | Phenix RealSpace Refine |
| Model resolution (Å)<br>FSC threshold | 3.4<br>0.5 |
| EMRinger score | 2.96 |
| CC (mask) | 0.81 |
| *Model composition*<br>Non-hydrogen atoms<br>Protein residues<br>Ligands | <br>8881<br>1151<br>0 |
| *Mean B factors ($Å^2$)*<br>Protein | <br>71.83 |
| *R.m.s. deviations*<br>Bond lengths (Å)<br>Bond angles (°) | <br>0.004<br>0.658 |
| *Validation*<br>MolProbity score<br>Clashscore<br>Poor rotamers (%) | <br>1.2<br>1.94<br>0.21 |
| *Ramachandran plot*<br>Favored (%)<br>Allowed (%)<br>Disallowed (%) | <br>96.37<br>3.63<br>0.00 |
| Cβ outliers (%) | 0.00 |
| CaBLAM outliers (%) | 2.26 |

**Supplementary Table 2**: Sequences of chimeric constructs and BRET constructs.

Legend: Hemagglutinin signal sequence, FLAG tag, PrC tag, residue swaps/additions

| Name | Sequence |
|---|---|
| FZD4 wild type | MKTIIALSYIFCLVFADYKDDDDKFGDEEERRCDPIRISMCQNLGYNVTKMPNL<br>VGHELQTDAELQLTTFTPLIQYGCSSQLQFFLCSVYVPMCTEKINIPIGPCGG<br>MCLSVKRRCEPVLKEFGFAWPESLNCSKFPPQNDHNHMCMEGPGDEEVPL<br>PHKTPIQPGEECHSVGTNSDQYIWVKRSLNCVLKCGYDAGLYSRSAKEFTDI<br>WMAVWASLCFISTAFTVLTFLIDSSRFSYPERPIIFLSMCYNIYSIAYIVRLTVGR<br>ERISCDFEEAAEPVLIQEGLKNTGCAIIFLLMYFFGMASSIWWVILTLTWFLAAG<br>LKWGHEAIEMHSSYFHIAAWAIPAVKTIVILIMRLVDADELTGLCYVGNQNLDAL<br>TGFVVAPLFTYLVIGTLFIAAGLVALFKIRSNLQKDGTKTDKLERLMVKIGVFSV<br>LYTVPATCVIACYFYEISNWALFRYSADDSNMAVEMLKIFMSLLVGITSGMWIW<br>SAKTLHTWQKCSNRLVNSGKVKREKRGNGWVKPGKGSETVV |
| FZD4 with ECL1 of FZD10 | MKTIIALSYIFCLVFADYKDDDDKFGDEEERRCDPIRISMCQNLGYNVTKMPNL<br>VGHELQTDAELQLTTFTPLIQYGCSSQLQFFLCSVYVPMCTEKINIPIGPCGG<br>MCLSVKRRCEPVLKEFGFAWPESLNCSKFPPQNDHNHMCMEGPGDEEVPL<br>PHKTPIQPGEECHSVGTNSDQYIWVKRSLNCVLKCGYDAGLYSRSAKEFTDI<br>WMAVWASLCFISTAFTVLTFLIDSSRFSYPERPIIFLSMCYNIYSIAYIVRLTVGA<br>ESIACDRDSGQLYVIQEGLESTGCAIIFLLMYFFGMASSIWWVILTLTWFLAAG<br>LKWGHEAIEMHSSYFHIAAWAIPAVKTIVILIMRLVDADELTGLCYVGNQNLDAL<br>TGFVVAPLFTYLVIGTLFIAAGLVALFKIRSNLQKDGTKTDKLERLMVKIGVFSV<br>LYTVPATCVIACYFYEISNWALFRYSADDSNMAVEMLKIFMSLLVGITSGMWIW<br>SAKTLHTWQKCSNRLVNSGKVKREKRGNGWVKPGKGSETVV |
| FZD4 with TM2 of FZD10 | MKTIIALSYIFCLVFADYKDDDDKFGDEEERRCDPIRISMCQNLGYNVTKMPNL<br>VGHELQTDAELQLTTFTPLIQYGCSSQLQFFLCSVYVPMCTEKINIPIGPCGG<br>MCLSVKRRCEPVLKEFGFAWPESLNCSKFPPQNDHNHMCMEGPGDEEVPL<br>PHKTPIQPGEECHSVGTNSDQYIWVKRSLNCVLKCGYDAGLYSRSAKEFTDI<br>WMAVWASLCFISTAFTVLTFLIDSSRFSYPERPIIFLSMCYCVYSVGYLIRLFAG<br>RERISCDFEEAAEPVLIQEGLKNTGCAIIFLLMYFFGMASSIWWVILTLTWFLAA<br>GLKWGHEAIEMHSSYFHIAAWAIPAVKTIVILIMRLVDADELTGLCYVGNQNLD<br>ALTGFVVAPLFTYLVIGTLFIAAGLVALFKIRSNLQKDGTKTDKLERLMVKIGVFS<br>VLYTVPATCVIACYFYEISNWALFRYSADDSNMAVEMLKIFMSLLVGITSGMWI<br>WSAKTLHTWQKCSNRLVNSGKVKREKRGNGWVKPGKGSETVV |
| FZD4 with TM3 of FZD10 | MKTIIALSYIFCLVFADYKDDDDKFGDEEERRCDPIRISMCQNLGYNVTKMPNL<br>VGHELQTDAELQLTTFTPLIQYGCSSQLQFFLCSVYVPMCTEKINIPIGPCGG<br>MCLSVKRRCEPVLKEFGFAWPESLNCSKFPPQNDHNHMCMEGPGDEEVPL<br>PHKTPIQPGEECHSVGTNSDQYIWVKRSLNCVLKCGYDAGLYSRSAKEFTDI<br>WMAVWASLCFISTAFTVLTFLIDSSRFSYPERPIIFLSMCYNIYSIAYIVRLTVGR<br>ERISCDFEEAAEPVLIQEGLKNTGCTLVFLVLYYFGMASSLWWVVLTLTWFLA<br>AGKKWGHEAIEMHSSYFHIAAWAIPAVKTIVILIMRLVDADELTGLCYVGNQNL<br>DALTGFVVAPLFTYLVIGTLFIAAGLVALFKIRSNLQKDGTKTDKLERLMVKIGV |

| | |
|---|---|
| FZD4 with CRD and Linker of FZD10 | MKTIIALSYIFCLVFADYKDDDDKISSMDMERPGDGKCQPIEIPMCKDIGYNMT<br>RMPNLMGHENQREAAIQLHEFAPLVEYGCHGHLRFFLCSLYAPMCTEQVSTP<br>IPACRVMCEQARLKCSPIMEQFNFKWPDSLDCRKLPNKNDPNYLCMEAPNN<br>GSDEPTRGSGLFPPLFRPQRPHSAQEHPLKDGGPGRGGCDNPGKFHHVEK<br>SASCAPLCGYDAGLYSRSAKEFTDIWMAVWASLCFISTAFTVLTFLIDSSRFSY<br>PERPIIFLSMCYNIYSIAYIVRLTVGRERISCDFEEAAEPVLIQEGLKNTGCAIIFL<br>LMYFFGMASSIWWVILTLTWFLAAGLKWGHEAIEMHSSYFHIAAWAIPAVKTIV<br>ILIMRLVDADELTGLCYVGNQNLDALTGFVVAPLFTYLVIGTLFIAAGLVALFKIR<br>SNLQKDGTKTDKLERLMVKIGVFSVLYTVPATCVIACYFYEISNWALFRYSADD<br>SNMAVEMLKIFMSLLVGITSGMWIWSAKTLHTWQKCSNRLVNSGKVKREKR<br>GNGWVKPGKGSETVV |
| FZD4 with Hinge of FZD10 | MKTIIALSYIFCLVFADYKDDDDKFGDEEERRCDPIRISMCQNLGYNVTKMPNL<br>VGHELQTDAELQLTTFTPLIQYGCSSQLQFFLCSVYVPMCTEKINIPIGPCGG<br>MCLSVKRRCEPVLKEFGFAWPESLNCSKFPPQNDHNHMCMEAPNNGSDEP<br>TRGSGLFPPLFRPQRPHSAQEHPLKDGGPGRGGCDNPGKFHHVEKSASCA<br>PLCGYDAGLYSRSAKEFTDIWMAVWASLCFISTAFTVLTFLIDSSRFSYPERPII<br>FLSMCYNIYSIAYIVRLTVGRERISCDFEEAAEPVLIQEGLKNTGCAIIFLLMYFF<br>GMASSIWWVILTLTWFLAAGLKWGHEAIEMHSSYFHIAAWAIPAVKTIVILIMRL<br>VDADELTGLCYVGNQNLDALTGFVVAPLFTYLVIGTLFIAAGLVALFKIRSNLQK<br>DGTKTDKLERLMVKIGVFSVLYTVPATCVIACYFYEISNWALFRYSADDSNMA<br>VEMLKIFMSLLVGITSGMWIWSAKTLHTWQKCSNRLVNSGKVKREKRGNGW<br>VKPGKGSETVV |
| FZD4 with Lower Hinge of FZD10 | MKTIIALSYIFCLVFADYKDDDDKFGDEEERRCDPIRISMCQNLGYNVTKMPNL<br>VGHELQTDAELQLTTFTPLIQYGCSSQLQFFLCSVYVPMCTEKINIPIGPCGG<br>MCLSVKRRCEPVLKEFGFAWPESLNCSKFPPQNDHNHMCMEGPGDEEVPL<br>PHKTPIQPGEECDNPGKFHHVEKSASCVLKCGYDAGLYSRSAKEFTDIWMAV<br>WASLCFISTAFTVLTFLIDSSRFSYPERPIIFLSMCYNIYSIAYIVRLTVGRERISC<br>DFEEAAEPVLIQEGLKNTGCAIIFLLMYFFGMASSIWWVILTLTWFLAAGLKWG<br>HEAIEMHSSYFHIAAWAIPAVKTIVILIMRLVDADELTGLCYVGNQNLDALTGFV<br>VAPLFTYLVIGTLFIAAGLVALFKIRSNLQKDGTKTDKLERLMVKIGVFSVLYTVP<br>ATCVIACYFYEISNWALFRYSADDSNMAVEMLKIFMSLLVGITSGMWIWSAKT<br>LHTWQKCSNRLVNSGKVKREKRGNGWVKPGKGSETVV |
| FZD4-NLuc | MKTIIALSYIFCLVFADYKDDDDKFGDEEERRCDPIRISMCQNLGYNVTKMPNL<br>VGHELQTDAELQLTTFTPLIQYGCSSQLQFFLCSVYVPMCTEKINIPIGPCGG<br>MCLSVKRRCEPVLKEFGFAWPESLNCSKFPPQNDHNHMCMEGPGDEEVPL<br>PHKTPIQPGEECHSVGTNSDQYIWVKRSLNCVLKCGYDAGLYSRSAKEFTDI<br>WMAVWASLCFISTAFTVLTFLIDSSRFSYPERPIIFLSMCYNIYSIAYIVRLTVGR<br>ERISCDFEEAAEPVLIQEGLKNTGCAIIFLLMYFFGMASSIWWVILTLTWFLAAG<br>LKWGHEAIEMHSSYFHIAAWAIPAVKTIVILIMRLVDADELTGLCYVGNQNLDAL<br>TGFVVAPLFTYLVIGTLFIAAGLVALFKIRSNLQKDGTKTDKLERLMVKIGVFSV<br>LYTVPATCVIACYFYEISNWALFRYSADDSNMAVEMLKIFMSLLVGITSGMWIW<br>SAKTLHTWQKCSNRLVNSGKVKREKRGNGWVKPGKGSETVVGGGSGGGS<br>GGGSVFTLEDFVGDWRQTAGYNLDQVLEQGGVSSLFQNLGVSVTPIQRIVLS<br>GENGLKIDIHVIIPYEGLSGDQMGQIEKIFKVVYPVDDHHFKVILHYGTLVIDGV<br>TPNMIDYFGRPYEGIAVFDGKKITVTGTLWNGNKIIDERLINPDGSLLFRVTING<br>VTGWRLCERILA |

| | |
|---|---|
| Tspan12-Venus | MEDQVDPRLIDGKAREDSVKCLRCLLYALNLLFWLMSISVLAVSAWMRDYLN<br>NVLTLTAETRVEEAVILTYFPVVHPVMIAVCCFLIIVGMLGYCGTVKRNLLLLAW<br>YFGSLLVIFCVELACGVWTYEQELMVPVQWSDMVTLKARMTNYGLPRYRWL<br>THAWNFFQREFKCCGVVYFTDWLEMTEMDWPPDSCCVREFPGCSKQAHQ<br>EDLSDLYQEGCGKKMYSFLRGTKQLQVLRFLGISIGVTQILAMILTITLLWALYY<br>DRREPGTDQMMSLKNDNSQHLSCPSVELLKPSLSRIFEHTSMANSFNTHFE<br>MEELGSGGSGVSKGEELFTGVVPILVELDGDVNGHKFSVSGEGEGDATYGK<br>LTLKLICTTGKLPVPWPTLVTTLGYGLQCFARYPDHMKQHDFFKSAMPEGYV<br>QERTIFFKDDGNYKTRAEVKFEGDTLVNRIELKGIDFKEDGNILGHKLEYNYNS<br>HNVYITADKQKNGIKANFKIRHNIEDGGVQLADHYQQNTPIGDGPVLLPDNHY<br>LSYQSKLSKDPNEKRDHMVLLEFVTAAGITLGMDELYK |
| CD151-Venus | MEDQVDPRLIDGKGEFNEKKTTCGTVCLKYLLFTYNCCFWLAGLAVMAVGIW<br>TLALKSDYISLLASGTYLATAYILVVAGTVVMVTGVLGCCATFKERRNLLRLYFIL<br>LLIIFLLEIIAGILAYAYYQQLNTELKENLKDTMTKRYHQPGHEAVTSAVDQLQQ<br>EFHCCGSNNSQDWRDSEWIRSQEAGGRVVPDSCCKTVVALCGQRDHASNI<br>YKVEGGCITKLETFIQEHLRVIGAVGIGIACVQVFGMIFTCCLYRSLKLEHYGS<br>GGSGVSKGEELFTGVVPILVELDGDVNGHKFSVSGEGEGDATYGKLTLKLICT<br>TGKLPVPWPTLVTTLGYGLQCFARYPDHMKQHDFFKSAMPEGYVQERTIFFK<br>DDGNYKTRAEVKFEGDTLVNRIELKGIDFKEDGNILGHKLEYNYNSHNVYITA<br>DKQKNGIKANFKIRHNIEDGGVQLADHYQQNTPIGDGPVLLPDNHYLSYQSK<br>LSKDPNEKRDHMVLLEFVTAAGITLGMDELYK |
